# The type 2 diabetes factor methylglyoxal mediates axon initial segment shortening and neuronal network activity changes

**DOI:** 10.1101/2021.05.10.443439

**Authors:** Ryan B. Griggs, Duc V.M. Nguyen, Leonid M. Yermakov, Jeneane M. Jaber, Jennae N. Shelby, Josef K. Steinbrunner, John A. Miller, Carlos Gonzalez-Islas, Peter Wenner, Keiichiro Susuki

**Author notes:** **Corresponding authors:** Ryan B. Griggs; or, Keiichiro Susuki, Department of Neuroscience, Cell Biology, and Physiology, Boonshoft School of Medicine, Wright State University, 3640 Colonel Glenn Highway, Dayton, Ohio 45435, USA. **Conflict of Interest Statement:** The authors declare no competing financial interests.

## Abstract

Recent evidence suggests that alteration of axon initial segment (AIS) geometry (i.e., length or position along the axon) contributes to CNS dysfunction in neurological diseases. For example, AIS length is shorter in the prefrontal cortex of type 2 diabetic mice with cognitive impairment. The key type 2 diabetes-related factor that alters AIS geometry is unknown. Here, we tested whether modifying the levels of insulin, glucose, or methylglyoxal, a reactive carbonyl species that is a metabolite of glucose, changes AIS geometry in mature cultures of dissociated postnatal mouse cortex using immunofluorescent imaging of the AIS proteins AnkyrinG and βIV spectrin. Neither insulin nor glucose modification appreciably altered AIS length. Elevation of methylglyoxal produced reversible AIS shortening without cell death. Multi-electrode array recordings revealed a biphasic effect of methylglyoxal on neuronal network activity: an immediate, transient ∼300% increase in spiking and bursting rates was followed by a ∼20% reduction from baseline at 3 h. AIS length was unchanged at 0.5 h or 3 h after adding methylglyoxal, whereas development of AIS shortening at 24 h was associated with restoration of spiking to baseline levels. Immunostaining for the excitatory neuron marker Ca^2+^/calmodulin-dependent protein kinase II alpha revealed AIS shortening in both excitatory and inhibitory neuron populations. This suggests that complex mechanisms maintain neuronal network operation after acute exposure to the disease metabolite methylglyoxal. Importantly, our results indicate that methylglyoxal could be a key mediator of AIS shortening during type 2 diabetes.

**Significance Statement:** Small changes in the structure of the axon initial segment affect neuronal function and may be a key mediator of neurological complications in various disease states. However, the specific disease factors that mediate structural changes at the axon initial segment are relatively unknown. This is the first study to show that increase of methylglyoxal is sufficient to reduce axon initial segment length and modulate neuronal network function. Methylglyoxal is a disease factor implicated in a wide variety of conditions including type 2 diabetes, Alzheimer’s disease, and aging. Thus, these findings could significantly impact the understanding of neurological complications in several disease states and are of broad pathophysiological relevance.

## 1. Introduction

The axon initial segment (AIS) is a specialized excitable domain within neurons that maintains neuronal polarity and regulates action potential generation (Rasband, 2010; Bender and Trussell, 2012). Voltage-gated sodium channels (Nav) are anchored at the AIS by sub-membranous cytoskeletal (e.g., βIV spectrin) and scaffolding (e.g., AnkyrinG) proteins (Nelson and Jenkins, 2017). Alteration of AIS constituents, such as mutations in ion channels or loss of protein complexes, is emerging as a key pathophysiology in a wide variety of neurological conditions in humans (reviewed in (Buffington and Rasband, 2011; Huang and Rasband, 2018)). In the brains of animals, subtle changes in AIS geometry (i.e., length or position along the axon) are reported in models of type 2 diabetes (Yermakov et al., 2018), Alzheimer’s disease (Marin et al., 2016), aging (Atapour and Rosa, 2017; Ding et al., 2018), stroke (Hinman et al., 2013), multiple sclerosis-related demyelination (Hamada and Kole, 2015; Radecki et al., 2018), and neuropathic pain (Shiers et al., 2018). This suggests altered AIS geometry could be a common mechanism of CNS dysfunction.

Patients with type 2 diabetes show slight decrement of cognitive performance across their lifespan and have increased risk of developing Alzheimer’s-like dementia (Biessels et al., 2014; Biessels and Despa, 2018). Some studies suggest that alteration of the AIS in brain regions important for cognition may change cognitive behavior in type 2 diabetes and related conditions. For example, concurrent cortical AIS shortening and cognitive impairment are present in type 2 diabetic db/db mice (Yermakov et al., 2018, 2019) and Alzheimer’s-like mice (Marin et al., 2016; Sri et al., 2019). In another study of mice, pharmacological treatment of neuropathic pain led to simultaneous reversal of cognitive impairment and AIS shortening in the infralimbic cortex (Shiers et al., 2018), supporting a link between AIS shortening in cortical neurons and cognition. To better understand the mechanisms underlying cognitive impairment in type 2 diabetes and related conditions, it is important to identify the factor(s) that initiate changes in AIS geometry in cortical neurons. In db/db mice, exercise treatment concomitantly ameliorated hyperglycemia and AIS shortening associated with type 2 diabetes (Yermakov et al., 2018), suggesting that type 2 diabetes-related factors such as high glucose or insulin resistance may alter AIS geometry. Another potential mediator of AIS shortening in type 2 diabetes is the reactive carbonyl species and glucose metabolite methylglyoxal. We recently reported that methylglyoxal disrupts AnkryinG at paranodes in CNS myelinated axons (Griggs et al., 2018), suggesting that methylglyoxal could affect AnkyrinG-containing AIS in cortical neurons. Importantly, disruption of methylglyoxal metabolism in humans is linked to the same neurological conditions as altered AIS geometry in animal models, including type 2 diabetes (Andersen et al., 2018), Alzheimer’s disease (Kuhla et al., 2007; Haddad et al., 2019), aging (Kuhla et al., 2006; Beeri et al., 2011; Srikanth et al., 2013), multiple sclerosis (Wetzels et al., 2019), and pain (Düll et al., 2019). The effect of methylglyoxal, glucose, or insulin on AIS geometry and the operation of neuronal networks is unknown.

To determine the key type 2 diabetes-related factor that leads to AIS shortening we utilized a reductionist approach. We exposed dense cultures of dissociated postnatal mouse cortex to modified levels of insulin, glucose, or methylglyoxal and quantified AIS geometry using AnkyrinG and βIV spectrin immunofluorescence. Since increase of methylglyoxal, but not insulin or glucose, was sufficient to shorten the AIS, we evaluated the temporal relationship between AIS length and neuronal network activity changes after methylglyoxal exposure. Finally, we tested the cell-type specificity of AIS shortening using the excitatory neuron marker Ca^2+^/calmodulin-dependent protein kinase II alpha (CaMKIIα). We are the first to uncover that short-term elevation of methylglyoxal, a metabolic factor implicated in many neurological disease states including type 2 diabetes, changes AIS geometry and transiently alters neuronal network activity.

## 2. Materials and Methods

### 2.1. Animals

Adult male and female C57BL/6J mice (The Jackson Laboratory, Bar Harbor, ME; RRID:IMSR_JAX:000664) aged 7-30 weeks were used for in-house breeding for postnatal cortical cultures. Mice were housed in Laboratory Animal Resources at Wright State University at 22-24°C and 20-60% humidity under 12-hour light / 12-hour dark conditions (lights on 7:00–19:00) with *ad libitum* access to food and water. Mice were provided wood-chip bedding, toilet paper roll cardboard tubes, shredded accordion paper, and thick cotton pads for enrichment and group housed except when single housing was required for breeding purposes. All animal procedures were approved by the Institutional Animal Care and Use Committee at Wright State University (Animal Use Protocol # 1113 & 1190) and carried out in accordance with the National Institutes of Health Office of Laboratory Animal Welfare Guide for the Care and Use of Laboratory Animals and the ARRIVE 2.0 guidelines (Percie du Sert et al., 2020).

### 2.2. Postnatal cortical cultures

Preparation and culturing of dissociated cells from postnatal mouse cortices was similar to as described previously (Beaudoin et al., 2012) utilizing the following materials: Neurobasal-A Medium (10888-022; Thermo), Neurobasal Plus Medium (A3582901, Thermo), Neurobasal-A without D-glucose or sodium pyruvate (A2477501, Thermo), B-27 Supplement (17504-044; Thermo), B-27 Plus Supplement (A3582801, Thermo), B-27 without Insulin (A1895601, Thermo), GlutaMAX Supplement (35050061; Thermo), Sodium Pyruvate (11360070, Thermo), Hibernate-A Medium (A1247501; Thermo Fisher Scientific), gentamycin (G1272; Sigma), papain (PDS Kit; LK003178; Worthington Biochemical Corporation, Lakewood, NJ), DNase I (90083; Thermo), fetal bovine serum (FBS; 26140-087; Thermo), poly-L-lysine (P2636; Sigma), coverslips pre-coated with poly-L-lysine (12mm #1.5; Biocoat 354085; Corning, Corning, NY or GG-12-1.5-PLL; Neuvitro Corporation, Vancouver, WA), Biosafety Cabinet Class II A2 (NU-540-500, Nuaire, Plymouth, MN), CO_2_ incubators (NU-5710, Nuaire).

#### Preparation of dissociated cortical cells

Plating Media (Neurobasal-A or Neurobasal Plus, 2 mM GlutaMAX, 2% B-27 or B-27 Plus, 5% FBS, 10 µg/mL gentamycin) and Growth Media (Neurobasal-A or Neurobasal Plus, 2 mM GlutaMAX, 2% B-27 or B-27 Plus) were prepared fresh, stored at 4°C, and used within 7 d. Papain (40 U/mL) was solubilized in Neurobasal-A or Hibernate-A Medium containing DNase I (40 U/mL) and pre-warmed at 37°C for 20 min. Whole brains were removed from decapitated mouse pups at postnatal day 0 (P0) and covered in ice-cold Hibernate-A. The left and right cortices were separated, meninges removed, minced into ∼2 mm^3^ pieces and transferred to a 15 mL centrifuge tube containing 1 mL ice-cold Hibernate-A. Cells were enzymatically and mechanically dissociated by adding 1 mL of pre-warmed, activated papain/DNase solution (20 U/mL final concentration), incubating in a 37°C bead bath for 20 min with gentle swirling every 5 min, followed by gentle trituration 3-5 times using a fire-polished 9 cm Pasteur pipet. Triturated solution containing cells and tissue debris was passed through a sterile 100 μm strainer, transferred to a 15 mL centrifuge tube, centrifuged 5 min at 250 x g, supernatant decanted, cell pellet resuspended in ∼1-2 mL Plating Media per cortical pair, cells counted using a Hemocytometer (0267110, Fisher), and concentration adjusted to 1 × 10^6^ cells per mL with additional Plating Media. Preparation of dissociated cells from both cortices from one postnatal mouse pup brain typically yielded circa 2 × 10^6^ cells; however, sometimes cells dissociated from multiple pup brains were combined into one tube if larger numbers of cells were needed for a given experiment, and this was deemed as one preparation.

#### Plating

Coverslips (12 mm), tissue culture dishes (35 mm), and multi-electrode arrays (MEAs) were pre-coated with poly-L-lysine diluted to 0.1 mg/mL in 0.1 M Borate buffer pH 8.5 and incubated >1 h to overnight at 37°C, washed with sterile water 2 times, allowed to completely dry in biosafety cabinet, and stored for up to 2 weeks at 4°C until use. Dissociated cortical cells from postnatal day 0 (P0) pups were pipetted onto pre-coated: 12 mm coverslips contained in 35 mm dishes (100,000 cells in 100 μL per coverslip, 3-4 coverslips per 35 mm dish) for immunofluorescence and cell viability assays, 35 mm dishes (1,000,000 cells in 1 mL) for Western blotting, or MEAs (50,000 cells in 50 μL) for neuronal network activity recordings, allowed to settle onto the substrate for at least 10 min, and Plating Media added up to a final volume of 2 mL (35 mm dishes) or 1 mL (MEAs). After overnight incubation, all the Plating Media was exchanged for Growth Media and cultures were maintained in Growth Media with 50% media changes every 3-4 days for coverslips/dishes or up to 7 days for MEAs where specialized membrane lids were used to prevent evaporation (ALAMEA-MEM5, ALA Scientific, Farmingdale, NY).

#### Characterization

Previous studies indicate that the AIS matures during the first week of culture and maximal levels of both AnkyrinG (Galiano et al., 2012) and Nav (Yang et al., 2007) protein are present circa day *in vitro* (DIV) 7. Therefore, drug exposure experiments were initiated on or after DIV8. The percentage of microtubule associated protein 2 (MAP2) positive neurons MAP2+ (78.2 ± 2.5 %) and glial fibrillary acidic protein (GFAP) positive astrocytes (21.8 ± 2.5 %) at DIV8 was similar to the percentage of MAP2+ (72.6 ± 3.4 %) and GFAP+ (27.4 ± 3.4 %) cells at DIV14 (n=12 coverslips from 4 preparations). This distribution of astrocytes (∼20%) is greater than the 6-8% previously reported in low density cortical cultures prepared from postnatal P0-1 (Beaudoin et al., 2012) or embryonic E17 (Belanger et al., 2011) mice. The percentage of MAP2+ neurons that were CaMKIIα+ (78.5 ± 1.07 %, n=12 coverslips from 3 preparations) was greater than CaMKIIα- (21.5 ± 1.07 %, n=12 coverslips from 3 preparations) at DIV14. This distribution of putative inhibitory cells (∼20%) agrees with previous mouse studies of GABA+ cells in cortical cultures (Beaudoin et al., 2012) or GAD67+ cells in hippocampal cultures (Prestigio et al., 2019). The density of cells in our cortical cultures (empirically determined to be ∼1,400 cells / mm^2^) is about three times the density of a previous study of AIS plasticity in hippocampal cultures where 45,000 cells were seeded onto 13 mm coverslips (calculated density of ∼ 440 cells / mm^2^) (Evans et al., 2013). This is notable, since a previous study in rat embryonic hippocampal cultures indicates that cell density alters neuronal excitability and AIS geometry (Guo et al., 2017).

### 2.3. Drug exposure

Dissociated P0 cortical cells were cultured on coverslips contained in 35 mm dishes for immunofluorescence or directly in 35 mm dishes for Western blotting. Cultures were exposed to insulin (A11382II; ThermoFisher, Waltham, MA), D-(+)-glucose (G8270; Sigma), D-mannitol (M4125; Sigma), or methylglyoxal (MG; 6.49 M stock; M0252; Sigma-Aldrich, St. Louis, MO) by dilution of the drug in Growth Media to 2x final concentration, syringe-filtration through 0.22 μm PES membranes (229746; CELLTREAT Scientific Products, Pepperell, MA), pre-warming to 37°C, and then adding 1 mL to 35 mm dishes containing 1 mL of conditioned medium for a 1x final concentration.

#### Insulin exposure

Insulin resistance was generated in our postnatal cortical cultures as previously described in embryonic mouse cortical cultures (Kim et al., 2011) with modification. Cultures were grown in Growth Media containing standard Neurobasal-A from DIV 0-10. Since the concentration of insulin in standard Neurobasal-A media is already supraphysiological (∼700 nM), we generated insulin resistance by first omitting insulin completely for 24 h at DIV 10-11 followed by 24 h exposure to standard Growth Media containing ∼700 nM insulin or Grown Media modified to contain 0, 20, or 100 nM insulin from DIV 11-12. We validated the generation of insulin resistance by challenging a subset of cultures grown on 35 mm dishes with 20 nM insulin for 15 min followed by quantification of the phosphorylated to non-phosphorylated Akt ratio (pAkt/Akt) using Western immunoblotting band density. Cultures used to quantify AIS length did not undergo the 15 min insulin challenge.

#### Glucose exposure

To model hyperglycemia in mature (DIV 10-11) cultures, coverslips were grown in standard culture conditions from DIV 0-10 and then exposed to media that was modified to contain 25 mM glucose (concentration in unmodified Neurobasal-A), 50 mM high glucose, or 25 mM normal glucose plus 25 mM mannitol. Mannitol was added to control for potential effect of hyperosmolality generated by doubling the amount of glucose in standard culture conditions (25 mM) to 50 mM high glucose.

#### Methylglyoxal exposure

Cultures were grown in standard Growth Media followed by methylglyoxal exposure initiated on DIV13 for Western immunoblotting and MEA analyses and DIV8-13 for immunofluorescence quantitation of AIS geometry or viability.

### 2.4. Western blotting of postnatal cortical cultures grown on 35 mm dishes

Dissociated cells from P0 mice cortices were grown on poly-L-lysine coated 35 mm tissue culture treated dishes as described above. Protein lysates for Western immunoblotting were prepared by rinsing dishes in ice-cold PBS twice then adding ice-cold lysis buffer (25 mM Tris HCl at pH 7.5, 150 mM NaCl, 1% Triton x100, 0.5% Deoxycholate, 0.1% SDS, 10 mM EDTA) containing protease inhibitor cocktail (P8340, Sigma) and phosphatase inhibitor cocktail (78428, Thermo). Cells were collected using a sterile cell scraper (08-100-241, Fisher) in 2 x 100 µL volumes of ice-cold lysis buffer, transferred to ice-cold 1.5 mL tubes, vortexed, incubated on ice for 10 min, then centrifuged at 18,000 x g for 10 min at 4°C in a Sorvall Legend Micro 21R centrifuge (Thermo). After centrifugation, supernatant was collected into fresh, ice-cold tubes and protein concentrations were measured using a Pierce BCA Protein Assay (232525, Thermo) or Coomassie Plus Bradford Assay (23236, Thermo). Samples (10 µg protein) were denatured at 95°C for 5 min in 4x Laemmli Sample Buffer (1610747, Bio-Rad) or Novex NuPAGE LDS Sample Buffer (NP0007, Thermo) with β-mercaptoethanol (1610710, Bio-Rad) or 10x reducing agent (DTT; B0009, Thermo), then run on either a 4–20% Mini-PROTEAN TGX stain-free gel (4568096, Bio-Rad) or on 4-12% gels using the Novex Bolt mini-gel system (NW0412B, B1000B, BT000614, Thermo). The gel was then transferred to nitrocellulose membrane, 0.45 µm pore size (Bio-Rad, Cat# 1620115). Membranes were blocked for 1 h in Pierce Protein-free (PBS) Blocking Buffer (37572, Thermo) for MG-H1 or 20 mM Tris, pH 8.0 and 0.05% (v/v) Tween 20 (TBST) containing 4% (w/v) milk for all other antigens. Membranes were incubated overnight at 4°C in primary antibody diluted (1:1000) in TBST with milk or protein-free blocking buffer. Primary antibody was washed 3 x 5 min in TBST followed by incubation of the membrane in horseradish peroxidase (HRP) conjugated secondary antibody (1:10,000 or 1:20,000; Jackson ImmunoResearch Laboratories) for 1 h at room temperature. Signals generated by Pierce ECL Plus Western Blotting Substrate (Thermo Scientific, Cat# 32132) were detected using a ChemiDoc MP Imaging System (Bio-Rad) or Azure 600 (Azure Biosystems, Dublin, CA, USA). Quantification of the band density was performed using ImageLab software (Bio-Rad). The densities of the bands of interest were normalized to the relative expression of GAPDH or Total Protein staining by TGX stain-free pre-cast gels (4568096, Bio-Rad), SYPRO Ruby (S4942, Sigma), or Azure Ponceau (10147-344, VWR).

### 2.5. Immunofluorescent analysis of postnatal cortical cultures grown on 12 mm coverslips

Immunostaining of postnatal cortical cultures was performed as follows. Custom humidity chambers were created by super gluing caps from 1.5 mL microcentrifuge tubes to the bottom of 100 mm Petri dishes and adding a piece of Kimwipe wetted with dH_2_O. After drug exposure, media was removed and dishes were rinsed three times in pre-warmed 1x PBS (137 mM NaCl, 2.7 mM KCl, 11.9 mM phosphate) then fixed in room temperature 4% paraformaldehyde (PBS, pH 7.4) for 20 min. After washing 3 x 5 min in PBS, coverslips were taken from the 35 mm dishes and dabbed on a Kimwipe to remove liquid then placed onto the microcentrifuge caps within the humidity chambers to complete the immunostaining process. Coverslips were blocked in PBSTGS (1x PBS, 0.1 % Triton X-100, 10% goat serum) for 30-60 min. Fifty microliters of primary antibody diluted in PBSTGS was added to each coverslip and then coverslips within the custom humidity chambers were incubated in primary antibody overnight at 4°C. After 3 x 5 min wash in PBSTGS, appropriate secondary antibodies were diluted in PBSTGS, added to the coverslips, and incubated in the dark at room temperature for 1 h. Coverslips were washed for 5 min each in PBSTGS, 0.01 M phosphate buffer, 0.005 M phosphate buffer, allowed to air dry, then mounted on non-coated slides using either KPL Mounting Medium (71-00-16; KPL, Inc.) or ProLong Gold Antifade Mountant without DAPI (36934; Thermo) or with DAPI (P36941; Thermo). In some instances, Hoechst was added during the PBSTGS wash step after secondary antibody incubation. After mounting onto slides, coverslips were secured using clear nail polish and stored at - 20°C until imaging analysis.

The primary antibodies used are listed in Table 1. AffiniPure minimally cross-reactive secondary antibodies used for immunofluorescent imaging were conjugated to Alexa Fluor (594, 488, 350) or AMCA (Jackson ImmunoResearch Laboratories, West Grove, PA). Secondary antibodies used for western immunoblotting were conjugated to horseradish peroxidase (Jackson ImmunoResearch Laboratories). Hoechst or DAPI were used to stain cell nuclei.

**Table 1.** Primary antibodies used for immunofluorescent imaging or Western immunoblotting.

| Antigen | Dilution | Species | Isotype | Manufacturer Information & RRID Citation |
| --- | --- | --- | --- | --- |
| Akt | 1:1000 | Rabbit | IgG | (Cell Signaling Technology Cat# 9272, RRID:AB_329827) |
| Phospho-Akt (Ser473) | 1:1000 | Rabbit | IgG | (Cell Signaling Technology Cat# 9271, RRID:AB_329825) |
| AnkyrinG | 1:400 | Mouse | IgG2a | (UC Davis/NIH NeuroMab Facility Cat# 75-146, RRID:AB_10673030) |
| βIV spectrin | 1:400 | Rabbit | IgG | (M.N. Rasband, Baylor College of Medicine, Texas, USA Cat# bIV SD, RRID:AB_2315634) |
| CaMKIIα | 1:400 | Mouse | IgG1 | (Thermo Fisher Scientific Cat# MA1-048, RRID:AB_325403) |
| GAPDH | 1:1000 | Mouse | IgG1 | (Enzo Life Sciences Cat# ADI-CSA-335-E, RRID:AB_2039148) |
| GFAP | 1:1000 | Mouse | IgG1 | (UC Davis/NIH NeuroMab Facility Cat# 75-240, RRID:AB_10672299) |
| MAP2 | 1:1000 | Chicken | IgY | (EnCor Biotechnology Cat# CPCA-MAP2, RRID:AB_2138173) |
| MG-H1 | 1:1000 | Rat | IgG | (T. Fleming & P. Nawroth, Heidelberg University, Germany; 1D2 tissue culture supernatant) |

Images of immunostained coverslips were obtained using a fluorescence microscope (Axio Observer Z1 with Apotome 2 fitted with AxioCam Mrm CCD camera; Carl Zeiss) and measurement of axon initial segment length was performed using the curve spline measurement tool in ZEN2.0 (Carl Zeiss) or a Matlab script adapted from previous studies (Grubb and Burrone, 2010; Yau et al., 2014). Our comparison of AIS length measured by eye versus by custom Matlab script yielded similar results, which is consistent with a strong correlation (Spearman r > 0.8) of AIS length measured by eye with AIS measurement using custom-written Matlab scripts as reported previously (Grubb and Burrone, 2010; Evans et al., 2015; Guo et al., 2017; Fruscione et al., 2018; Zhang et al., 2019). Measurement of AIS location (distance from the edge of the soma to the start of the AIS) was performed as described previously (Guo et al., 2017). Only continuous, non-overlapping AISs originating from a MAP2+ cell were quantified by an observer blinded to experimental treatment.

### 2.6. Cell viability assay

Cell viability was determined using a commercially available dual-wavelength fluorescent kit (LIVE/DEAD Viability/Cytotoxicity Assay Kit; L3224; Invitrogen). Live cells are detected by intracellular esterase activity using the dye calcein-AM (∼495/515 nm ex/em; green). Dead cells are detected by a large increase in fluorescence of the dye ethidium homodimer-1 when bound to nucleic acids, which only occurs if the plasma membrane is compromised (∼495/635 nm ex/em; red). Optimal dilution (1:2000 for each dye) of calcein-AM (4 mM in DMSO) and EthD-1 (2 mM in DMSO:H_2_O 1:4) stocks was empirically determined by comparing signal:noise ratios in the green and red channels. We validated the live/dead assay by exposing mature cortical cultures to either 1x PBS or 100% methanol for 15 min and confirming methanol-induced cell death. After drug exposure, coverslips were rinsed 3x in 1x PBS and incubated in 80 µL of calcein-AM/EthD-1/PBS solution in the dark at room temperature for 15 min. Excess solution was removed and coverslips were mounted in 5 µL calcein-AM/EthD-1/PBS solution, sealed using clear nail polish to minimize evaporation, and immediately imaged to avoid signal decay of calcein-AM fluorescence. The number of cells displaying green or red fluorescence was quantified and data is presented as percentage of # live cells counted divided by total cells counted (# live cells + # dead cells) normalized to media control.

### 2.7. Multielectrode array recording of neuronal network activity

Multielectrode array (MEA) recordings were performed as previously described (Wagenaar et al., 2006; Hales et al., 2010; Fong et al., 2015) using the MEA2100-60-System (Multi Channel Systems, Germany) that includes MEA2100-HS60 headstage with integrated heating element, MCS-interface board 3.0 multiboot, Multi Channel Suite acquisition and analysis software, and temperature controller. MEAs (60MEA200/30iR-Ti-gr; Multi-Channel Systems, Reutlingen, Germany) were pre-coated with poly-L-lysine as described above, a 25 μL drop of 40 µg/mL laminin (L2020, Sigma) was carefully pipetted directly onto the center of the electrode array, and MEAs were incubated at 37°C during preparation of postnatal cortical cells. Prior to plating the cells, laminin was incompletely aspirated leaving a wetted area that localizes dispersion of the cell suspension on top of the electrode array. MEAs were maintained by 50% media changes every 3-7 d with fresh pre-warmed Growth Media using specialized gas-permeable membrane ALA lids to prevent evaporation that can result in increased tonicity and toxicity (Potter and DeMarse, 2001). MEAs remained in the incubator during the experiment except during the 15 min recording period at each timepoint. Recordings were performed with the MEA2100-60-System on the benchtop in ambient air with headstage heated to 37°C. The ALA lids prevented contamination during repeated recordings across several days. Since we observed that 50% media changes altered spiking activity within the first several hours, experimental recordings were performed the day after media changes. Vehicle control (2 µL Growth Media) or methylglyoxal (2 μL of 500x final concentration) were added to MEAs containing 1 mL conditioned media resulting in a 1x final concentration. The pH was similar before and after addition of either media or methylglyoxal. Recordings were digitized at 20,000 Hz. Threshold for spike detection was set at mean noise ± 5 standard deviations on a per channel basis as previously reported (Charkhkar et al., 2015). Multi Channel Suite (Multi Channel Systems) and Neuroexplorer 5.0 (Nex Technologies, Colorado Springs, CO) were used to create raster plots and extract features of network spiking and bursting.

MEA-wide network spike frequency was obtained by dividing the recording duration (15 minutes) into the total amount of spikes from 59 channels (reference channel was omitted) (Wagenaar et al., 2006; Fong et al., 2015). Network bursts were extracted using the MCS Analyzer Burst Analyzer tool with settings of 50 ms maximum interval to start/end burst, 100 ms minimum interval between bursts, 50 ms minimum duration of burst, at least 5 spikes within a burst, and at least 15 channels participating in the network burst. MEA-wide network bursting, defined as synchronized spiking activity across all or most of the recording channels, was consistently seen in all MEA preparations. The variance of network spike frequency in the MEAs at baseline was relatively large; therefore, results are presented as percentage of baseline network spike frequency within each MEA. To account for heterogeneity of network function between culture preparations as previously reported (Wagenaar et al., 2006), we utilized replicate (sister cultures) MEAs from multiple preparations.

### 2.8. Experimental Design and Statistical Analysis

Comparison of the means between two groups was performed using unpaired, homoscedastic t-test. Effect of insulin, glucose, or methylglyoxal concentration was analyzed using one-way ANOVA and/or linear regression. An interaction of time and drug was determined using two-way ANOVA. Effect of a single methylglyoxal dose over time was determined by repeated measures ANOVA. Multiple comparison post-hoc tests are listed in the results text. An alpha value of α=0.05 was used to determine statistical significance. All data were analyzed using Prism 9.0 (GraphPad, La Jolla, CA) or SAS 9.4 (SAS Institute, Inc., Cary, NC) and are presented as cumulative frequency distributions or using scatter plots or bar graphs with mean ± SEM.

## 3. Results

### 3.1. Insulin sensitivity manipulation does not affect AIS length

Type 2 diabetes is canonically characterized by insulin resistance and elevated glucose. First, we tested whether altering insulin sensitivity in postnatal cortical cultures affects AIS length. We utilized a protocol previously established to produce insulin resistance after 24 h exposure to insulin in embryonic mouse primary cortical cultures (Kim et al., 2011). Previous studies also suggest that a 24 h period of time should be sufficient to observe changes in AIS geometry in cultured CNS neurons (Grubb and Burrone, 2010; Chand et al., 2015). As shown in the experimental design (Fig. 1A), we varied the insulin concentration for 24 h in mature cultures and then validated insulin resistance via Akt immunoblotting and quantified AIS length by immunostaining for AnkryinG, the master organizer of the AIS (Rasband, 2010). Western blot image and quantification of phosphorylated Akt and total Akt band density revealed that increasing the concentration of insulin for 24 h led to a lower pAkt/Akt ratio when challenged with insulin for 15 min (Fig. 1B,C). Decreased phosphorylation of Akt is indicative of insulin resistance. A representative image of the AIS (AnkyrinG) and the somatodendritic domain (MAP2) in untreated cultures is shown (Fig. 1D). AIS length was similar after 24 h exposure to 700 nM (28.61 ± 0.28 µm, n=416 AIS from 3 preparations), 0 nM (28.41 ± 0.29 µm, n=396 AIS from 3 preparations), or 100 nM (28.20 ± 0.34 µm, n=293 AIS from 3 preparations) insulin (insulin concentration: *F*(2, 1102) = 0.45, *p* = 0.64; Tukey) (Fig. 1E). Cumulative frequency plot shows overlapping distributions of AIS lengths after variable insulin exposure (Fig. 1F), indicating that no changes in AIS length are observed in the entire population of cortical neurons.

**Figure 1.**
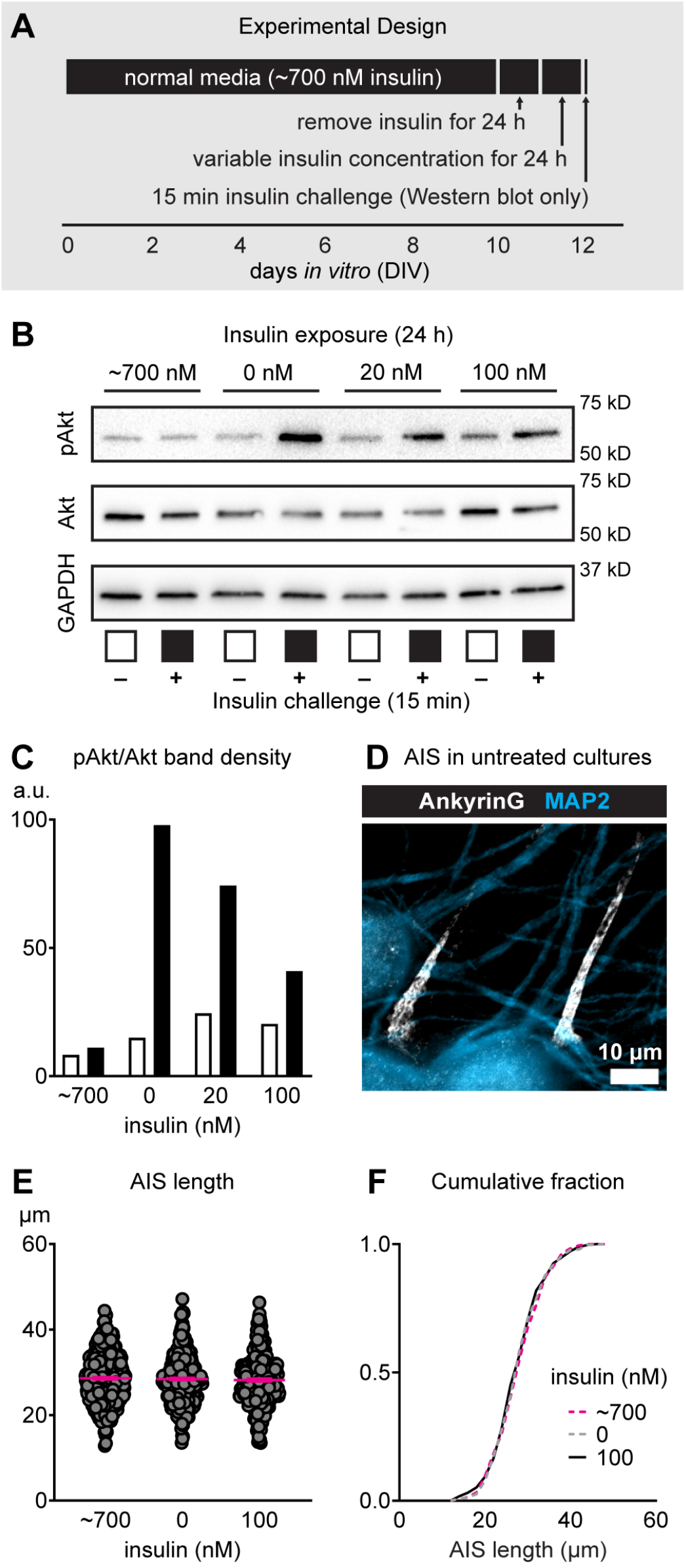
Effect of altering insulin on AIS length in mature postnatal mouse cortical cultures. (A) Schematic showing the experimental design and timeline of variable insulin exposure to induce insulin resistance (revealed by Western blotting after 15 min challenge to insulin) in culture. (B-C) Western immunoblotting results showing phosphorylation status of Akt after inducing acute insulin resistance. White bars denote absence of 15 min insulin challenge and black bars denote presence of 15 min insulin challenge that was used to reveal insulin sensitivity status. (D) Representative image of mature postnatal cortical cultures at DIV10 showing MAP2 (somatodendritic domain) and AnkyrinG (AIS). Scale bar = 10 µm. (E) AIS length after inducing conditions of complete insulin resistance (normal media containing ∼700 nM insulin), no insulin resistance (0 nM insulin), and moderate insulin resistance (100 nM insulin). (F) Cumulative fractional distribution of AIS lengths after 24 h exposure to ∼700, 0, or 100 nM insulin.

### 3.2. Glucose does not appreciably change AIS length

Next, we tested if elevating glucose levels in mature postnatal cortical cultures changes AIS length. AIS length was significantly altered (main effect of treatment: *F*(2, 844) = 3.56, *p* = 0.03; Tukey) after 24 h exposure to 25 mM glucose (30.92 ± 0.37 µm, n=291 AIS from 3 preparations), 50 mM high glucose (29.66 ± 0.37 µm, n=236 AIS from 3 preparations), or 25 mM glucose plus 25 mM mannitol (30.81 ± 0.32 µm, n=320 AIS from 3 preparations) (Fig. 2A). High glucose (50 mM) shortened the AIS by ∼4% compared to standard culture conditions containing 25 mM glucose (*p* = 0.04, Tukey). However, AIS length after 50 mM high glucose or osmotic control of 25 mM glucose plus 25 mM mannitol was similar (*p* = 0.06, Tukey), suggesting that high osmolality may play a role in the ∼4% AIS shortening produced by high glucose alone. Alternatively, diabetic hyperglycemia leads to accumulation of methylglyoxal in rodents and humans *in vivo* (Bierhaus et al., 2012; Sveen et al., 2013; Griggs et al., 2016, 2019; Huang et al., 2016) and increasing the glucose concentration in culture media from 5 to 20 mM results in a 2-fold intracellular and 5-fold extracellular increase of methylglyoxal *in vitro* (Irshad et al., 2019). Thus, we speculated that accumulation of methylglyoxal caused by the substantial increase of glucose may contribute to mild AIS shortening.

**Figure 2.**
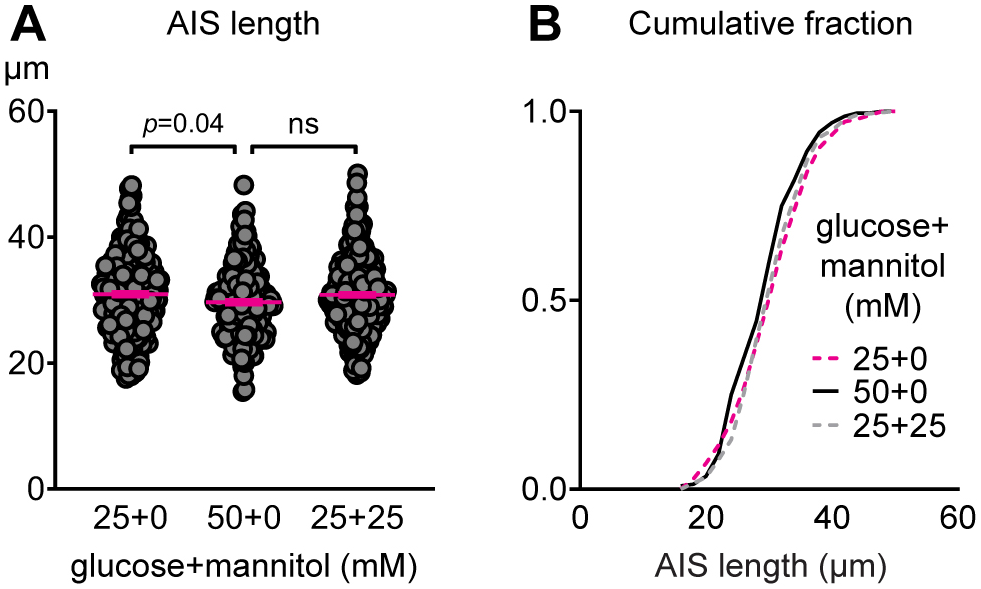
Effect of altering glucose on AIS length. (A) AIS length after 24 h exposure to 25 mM glucose, 50 mM glucose, or 25 mM glucose plus 25 mM mannitol as osmotic control in mature cultures (DIV 10-11). (B) Cumulative fractional distribution of AIS lengths after 24 h exposure to 25 mM glucose, 50 mM glucose, or 25 mM glucose plus 25 mM mannitol.

### 3.3. Methylglyoxal exposure increases cellular methylglyoxal-derived MG-H1

To test the hypothesis that accumulation of methylglyoxal disrupts AIS geometry, we first needed to determine the dose and duration of methylglyoxal exposure required to disrupt cellular methylglyoxal metabolism. Pathophysiological increase of methylglyoxal leads to accumulation of advanced glycation end-products (AGE). Upwards of 90% of MG-AGEs *in vivo* are methylglyoxal-derived hydroimidazolone (MG-H1) (Rabbani and Thornalley, 2012), and thus MG-H1 provides a suitable marker for disrupted methylglyoxal metabolism. Reliable estimates of methylglyoxal concentration in the nervous system are 1.2 µM (Eberhardt et al., 2012) or 40 μM (Liu et al., 2017) in dorsal root ganglia, 0.3-1.5 µM (Kurz et al., 2011) or 5 μM in young adult mouse brain (Distler et al., 2012), and 10 µM in the cerebrospinal fluid of healthy control middle-aged humans (Kuhla et al., 2005). Thus, we tested if exposing postnatal cortical cultures to 0, 1, 10, or 100 µM methylglyoxal for 24 h was sufficient to elevate cellular methylglyoxal to pathophysiological levels resulting in accumulation of MG-H1. MG-H1 and Total Protein in immunoblots of postnatal cortical culture cell lysates are shown (Fig. 3A). MG-H1 relative band density (normalized to total protein lane volume) was increased by 100 µM methylglyoxal (*p* = 0.001) but not 1 or 10 µM (*F* (3, 8) = 16.08; *p* = 0.0009; Dunnett; n=3 preparations per dose) (Fig. 3B). Our results indicate that an approximate 10-to 100-fold increase of normal methylglyoxal concentration in postnatal cortical cultures is sufficient to acutely disrupt cellular methylglyoxal metabolism.

**Figure 3.**
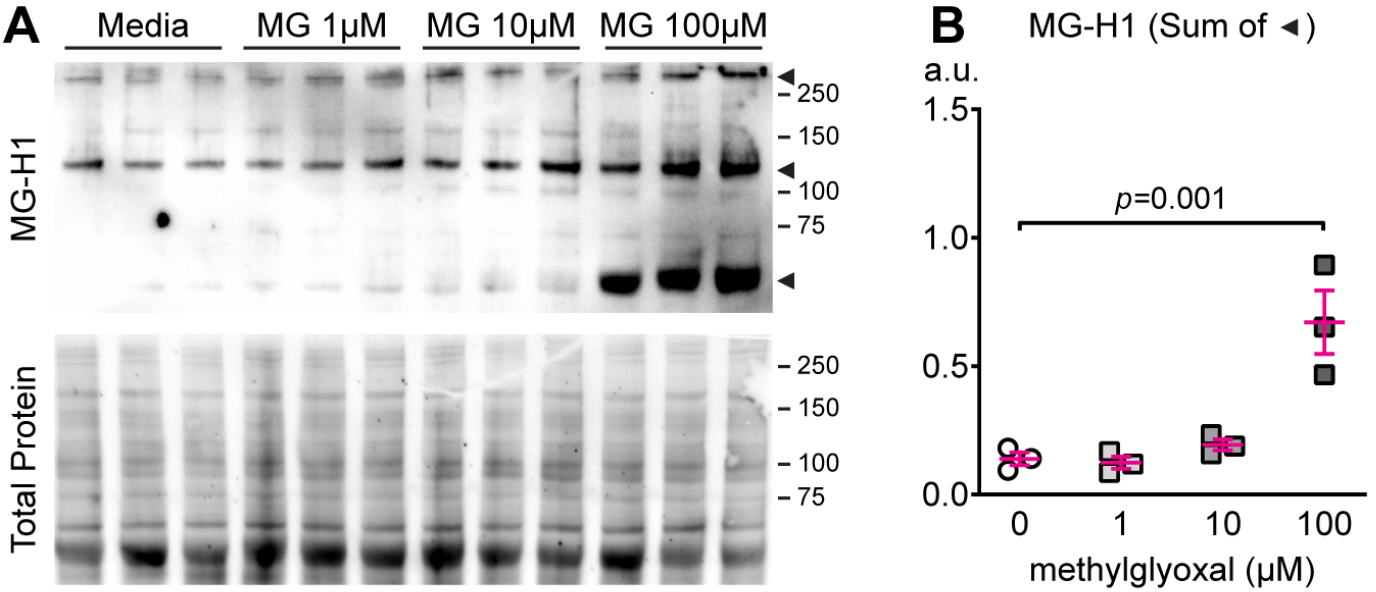
Methylglyoxal increases cellular methylglyoxal-derived MG-H1. (A) Western immunoblot image of cultures exposed to methylglyoxal (MG; 0-100 μM; 24 h; n=3) showing MG-H1 (unpurified rat anti-MG-H1 clone 1D2 antibody) and Total Protein (SYPRO Ruby). (B) Quantification of MG-H1 immunoblot density normalized to SYPRO Ruby total protein stain. The sum of band density for the three bands indicated by arrowheads was used for analysis.

### 3.4. Characterization of AIS geometry after methylglyoxal exposure

To test if methylglyoxal shortens the AIS, we exposed mature neuronal networks to 0-100 µM methylglyoxal for 24 h. Representative images of AnkyrinG and microtubule associated protein 2 (MAP2; somatodendritic marker) show a shorter AIS length after 100 µM methylglyoxal exposure for 24 h (Fig. 4A). AIS length after 24 h exposure to 0 µM (media control; 28.79 ± 0.28 µm, n = 598 AIS from 6 preparations), 1 µM (28.39 ± 0.36 µm, n = 326 AIS from 3 preparations), 10 µM (27.26 ± 0.29 µm, n = 489 AIS from 3 preparations), or 100 µM (25.48 ± 0.26 µm, n = 522 AIS from 4 preparations) methylglyoxal indicate a dose-dependent inverse relationship between concentration of methylglyoxal and AIS length (slope = -1.47, R^2^ = 0.85, *p* < 0.0001; linear regression analysis of means) with significant AIS shortening by 10 µM (*p* = 0.0004, 5.3% reduction) and 100 µM (*p* < 0.0001, 11.5% reduction) methylglyoxal compared to media only (dose: *F*(4, 2159) = 63.60, *p* < 0.0001; Dunnet) (Fig. 4B). This 11.5% AIS shortening induced by 100 µM methylglyoxal is about half of the 25% shortening induced by patterned optogenetic stimulation or high KCl (Evans et al., 2015) and similar to 8-16% decrease in AIS length in cortical neurons of db/db mice (Yermakov et al., 2018). AIS shortening by 100 µM methylglyoxal was generalized across the population of neurons, as indicated by a leftward shift in the cumulative frequency distribution (Fig. 4C). Taken together, our results indicate that methylglyoxal, but not insulin resistance or high glucose alone, could be the key initiator of AIS shortening during type 2 diabetes.

**Figure 4.**
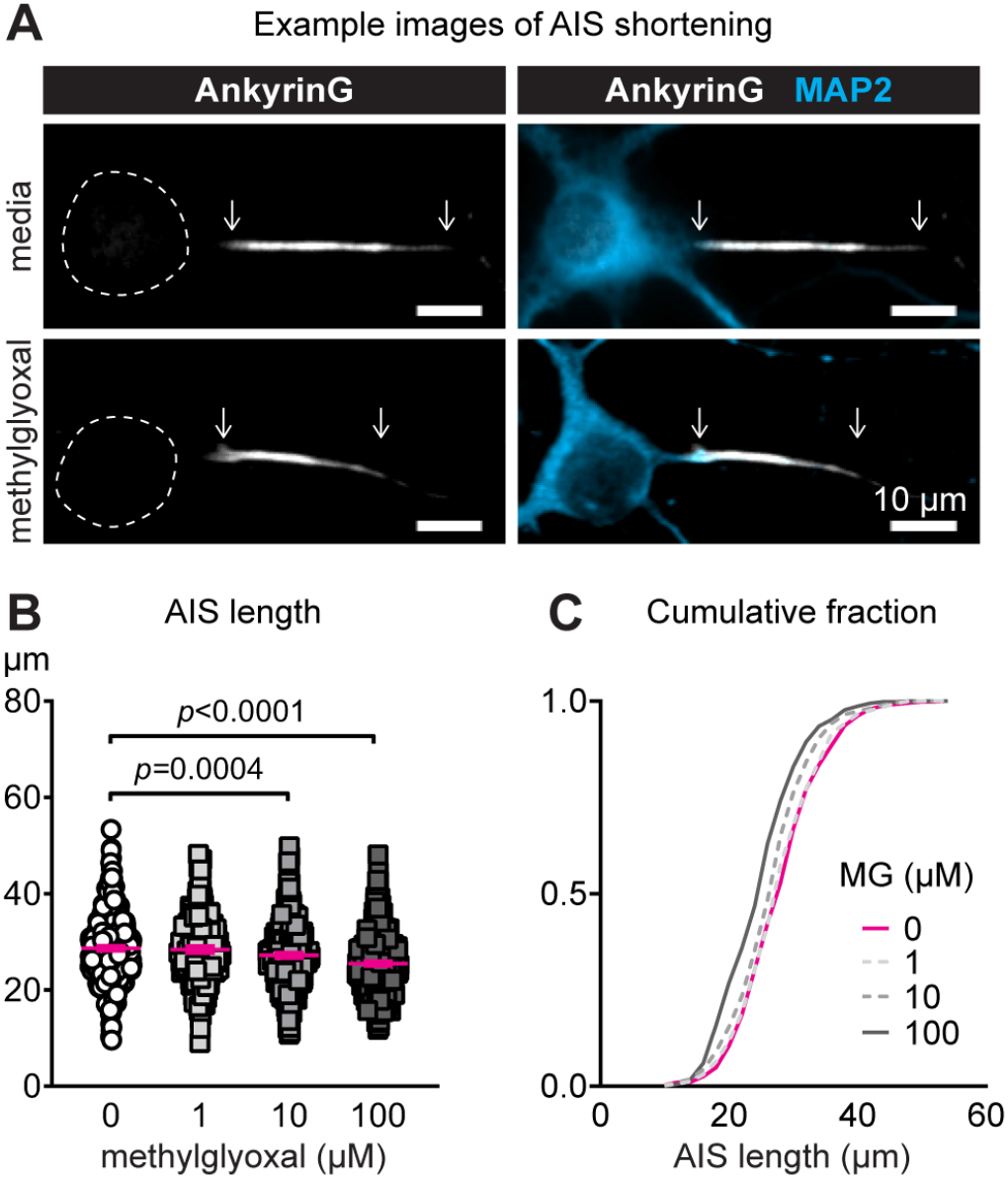
Methylglyoxal reduces AIS length in mature postnatal mouse cortical cultures. (A) AnkyrinG (AIS protein) and MAP2 (somatodendritic protein) immunofluorescence after 24 h exposure to media or 100 µM methylglyoxal at DIV10-11. Dashed lines indicate the neuronal soma. Scale bar = 10 µm. (B) AIS length (AnkyrinG immunostaining) after 24 h exposure to various concentrations of methylglyoxal (0-100 µM). (C) Cumulative fractional distribution of AIS lengths after 24 h exposure to 0-100 µM methylglyoxal (MG).

In addition to AIS length changes, AIS relocation (change in the distance from the soma to AIS start point along the axon) is implicated in experimental demyelination (Hamada and Kole, 2015), epilepsy (Harty et al., 2013), and as a mechanism of activity-dependent plasticity that alters intrinsic neuronal function (Grubb and Burrone, 2010; Evans et al., 2013, 2017). Therefore, we tested whether exposure to 100 µM methylglyoxal for 24 h relocated the AIS. AIS start location after 24 h exposure to media (5.23 ± 0.24 µm, n = 194 AIS from 3 preparations) or 100 µM methylglyoxal (4.70 ± 0.17, n = 246 AIS from 3 preparations) was similar (*p* = 0.07, unpaired homoscedastic two-tailed t-test). This is consistent with no change in the AIS start location in db/db prefrontal cortex (Yermakov et al., 2018).

### 3.5. Methylglyoxal-evoked AIS shortening is not associated with cell death and is reversible

To rule out the possibility that AIS shortening by methylglyoxal is due to cellular injury and/or loss of the AIS structure we assessed cell viability after methylglyoxal exposure. Live/dead analysis of exposure to methanol, a positive control, or 24 h exposure to a supraphysiological dose of 10 mM methylglyoxal showed substantial cell death (Fig. 5A). However, 100 µM methylglyoxal did not affect cell viability after 24 h exposure (*p* = 0.53; unpaired homoscedastic t-test; n=8 coverslips from 3 preparations) compared to media (Fig. 5B). Our results are in agreement with the absence of cell death in cultured embryonic (E17) mouse cortical neurons at DIV14 (Belanger et al., 2011), DRG neurons (Radu et al., 2012), or SH-SY5Y cells (Nishimoto et al., 2017) after *in vitro* exposure to 100 µM methylglyoxal for 24 h. Similarly *in vivo*, intraperitoneal injection of methylglyoxal increased the concentration of methylglyoxal in the brain without cytotoxicity in the hippocampus (Distler et al., 2012). In type 2 diabetic db/db mice at 10-11 weeks of age, there is elevated MG-H1 in serum (Griggs et al., 2019) and AIS shortening without apoptotic cell death in the prefrontal cortex (Yermakov et al., 2018).

**Figure 5.**
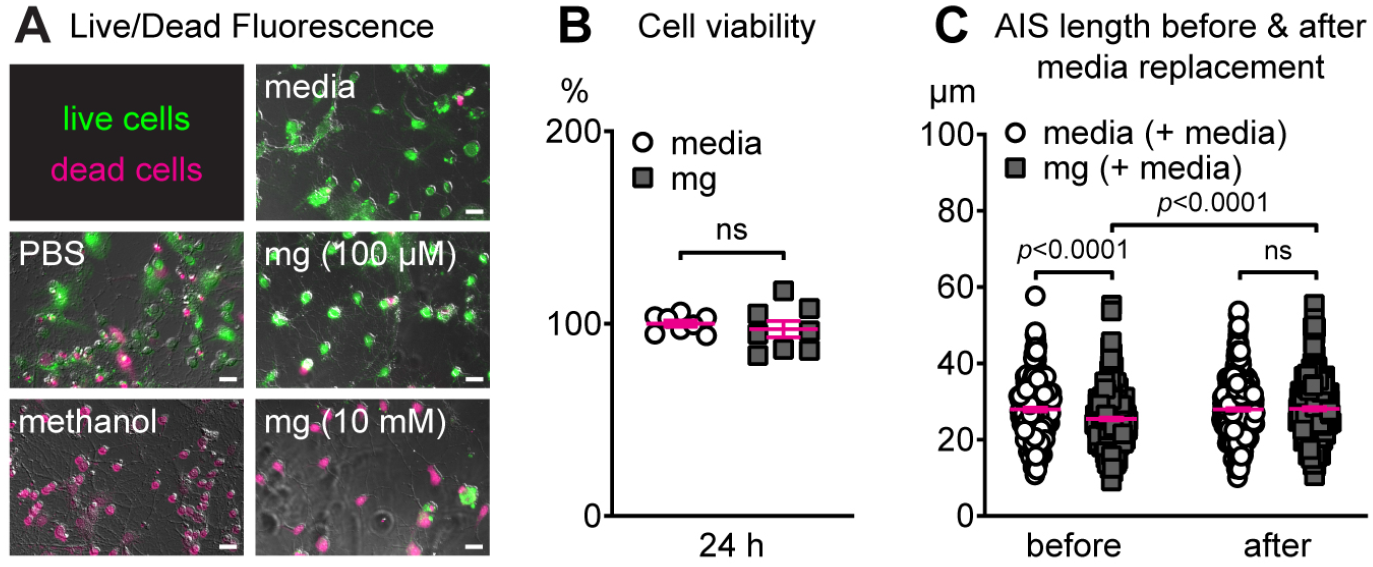
AIS shortening by methylglyoxal does not alter cell viability and is reversible. (A) Representative images of live/dead assay results after PBS (control) or methanol (to induce cell death) exposure for 15 min and after 24 h exposure to media, 100 µM methylglyoxal (mg), or 10 mM methylglyoxal (mg). Scale bar = 20 µm. (B) Cell viability (live/dead assay) results after 24 or 72 h exposure to media or 100 µM methylglyoxal (mg). (C) AIS length after an initial exposure to media or 100 µM methylglyoxal (mg) for 24 h (before), followed by replacement with conditioned media and a 24 h recovery period (after).

A previous study of AIS plasticity in embryonic rat hippocampal neurons indicates that AIS shortening initiated by high KCl is recoverable when cultures are returned to conditioned media for 24 h (Evans et al., 2015). Thus, we tested if media replacement would lead to recovery of AIS shortening after an initial 24 h exposure to 100 µM methylglyoxal (Fig. 5C). There was a significant interaction of AIS length between media or methylglyoxal exposure and before or after media replacement (drug by time interaction: *F*(1, 1496) = 10.48, *p* = 0.001; Tukey). Before media replacement, methylglyoxal AIS length (25.46 ± 0.35 µm, n=383 AIS from 3 preparations) was shorter compared to media (27.9 ± 0.46 µm, n=274 AIS from 3 preparations) (*p* = 0.0002). Twenty-four hours after media replacement, AIS length was similar in cultures initially exposed to media (media + media, 27.92 ± 0.33 µm, n=499 from 3 preparations) or methylglyoxal (methylglyoxal + media, 28.10 ± 0.40 µm, n=343 from 3 preparations) (*p* > 0.99). Importantly, since 24 h exposure to 100, but not 1 or 10, µM methylglyoxal was sufficient to elevate cellular methylglyoxal (Fig. 3) and shorten the AIS (Fig. 4) in a reversible manner without cell death (Fig. 5), this dose was used to further characterize the effect of methylglyoxal on the AIS and neuronal network activity.

### 3.6. Immediate effect of methylglyoxal on neuronal network activity

Burgeoning research suggests activity-dependent structural plasticity of AIS geometry is a mechanism of fine-tuning neuronal excitability (Yamada and Kuba, 2016; Jamann et al., 2018). For example, depolarization by high KCl induces AIS shortening at 1 h (Jamann et al., 2021), 3 h (Evans et al., 2015), or 24 h (Chand et al., 2015). Methylglyoxal acutely depolarizes neocortical pyramidal neurons in rat somatosensory cortex slices (de Arriba et al., 2006), suggesting the potential for depolarization-induced AIS plasticity in our postnatal cortical cultures. As a first step to elucidating whether AIS shortening by methylglyoxal is activity-dependent, we determined the immediate functional effect of methylglyoxal on neuronal network activity using multielectrode array (MEA) recordings. Characterization of MEA network activity prior to drug exposure revealed that MEA-wide network spike frequency increased from DIV7 (44.5 ± 4.8 Hz) to DIV13 (230.6 ± 10.9 Hz) (*p* < 0.0001; n=9 MEAs from 3 preparations). Example sixty second raster plots from DIV13 show that methylglyoxal immediately increases network activity, followed by a short period of quiescence, and restoration to baseline activity within 15 minutes (Fig. 6A). Methylglyoxal altered network spike rate compared to media control (drug x time interaction: *F* (4, 52) = 31.16; *p* < 0.0001; Sidak) (Fig. 6B). Methylglyoxal increased network spike rate (363.0 ± 33.5 %) compared to media control (117.7 ± 7.5 %) during the first minute by ∼300%. At 3 min, methylglyoxal reduced network spike rate (40.9 ± 9.5 %) to ∼50% of media control (89.3 ± 9.6 %). At 15 min, network spike rates were similar to baseline after either methylglyoxal (*p* = 0.99) or media (*p* = 0.73) exposure. Similarly, methylglyoxal increased the network burst rate (# of MEA-wide network bursts per minute) immediately after its addition with restoration of baseline bursting at 15 min (drug x time interaction: *F* (4, 52) = 24.45; *p* < 0.0001; Sidak) (Fig. 6C). Acute increase of network activity by methylglyoxal could lead to rapid (within 1-3 h) activity-dependent AIS shortening, so in the next experiments we characterized the time course of AIS shortening and network activity changes.

**Figure 6.**
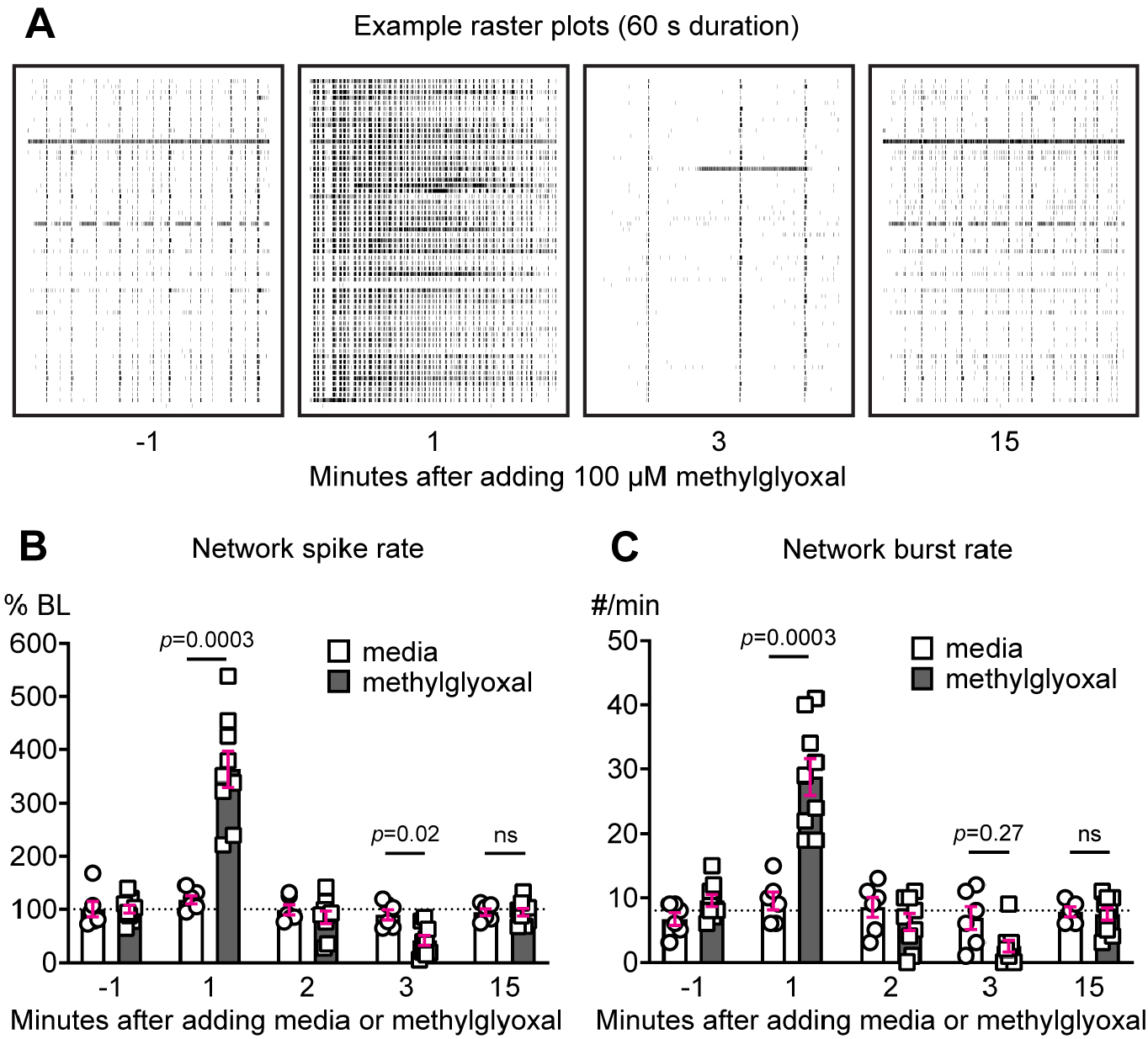
Methylglyoxal immediately increases neuronal network activity. (A) Representative raster plots (60 s duration) of MEA recordings showing spike activity (60 electrodes, one per row) during 1 min intervals before and after exposure to 100 µM methylglyoxal. (B) Quantification of network spike rate (normalized as percent of -1 min baseline) during 1 min intervals before and after exposure to media or 100 µM methylglyoxal. Dashed line indicates baseline. (C) Quantification of network burst rate (MEA-wide bursts per minute) during 1 min intervals before and after exposure to media or 100 µM methylglyoxal. Dashed line indicates baseline.

### 3.7. Time course of changes in AIS length and neuronal network activity

At the single cell level, AIS shortening dampens multiple spike firing and increases firing threshold *in vitro* (Evans et al., 2015) and *in vivo* (Jamann et al., 2021). *In silico*, AIS shortening is associated with increase of interspike interval (Baalman et al., 2013), voltage threshold (Baalman et al., 2013; Goethals and Brette, 2020), and action potential acceleration (Vascak et al., 2017) in single neurons. However, surprisingly little is known about how these changes in AIS length affect neuronal network operation. Thus, to test if there is an association between AIS shortening and altered network function we assessed AIS length and MEA spiking and bursting at 0.5, 3, and 24 h after addition of media or 100 µM methylglyoxal. There was a significant drug by time interaction in AIS length (*F* (2, 783) = 7.7, *p* = 0.0005; n=59-212 AIS from 1 preparation; Sidak). Example images of AnkryinG immunostaining show a shorter AIS at 24 h but not 0.5 h or 3 h after methylglyoxal exposure (Fig. 7A). AIS length after media exposure was similar at all timepoints (Fig. 7B), whereas methylglyoxal reduced AIS length at 24 h (Fig. 7C). Example raster plots of neuronal network activity are shown at baseline and at 3 and 24 h after exposure to 100 µM methylglyoxal (Fig. 7D). Control exposure to media alone did not alter network spike frequency (repeated measures across time: *F*(2, 10) = 0.74, *p* = 0.5; n=6 MEAs from 2 preparations) (Fig. 7E). Methylglyoxal altered network spike rate (repeated measures across time: *F*(2,16) = 20.31, *p* = 0.001; n=9 MEAs from 3 preparations), with reduced spiking at 3 h that returned to baseline level at 24 h (Fig. 7F). Media control did not alter network burst spikes (# of spikes within network bursts; Fig. 7G) or network burst duration (Fig. 7H). Network burst rate (# of network bursts per min) in media control MEAs slightly increased during the experiment (Fig. 7I). Network burst spike rate (# of spikes per second within a network burst) was lower at 3 h and unchanged at 24 h by media control treatment (Fig. 7J). Methylglyoxal altered network burst spikes (Fig. 7K) and network burst duration (Fig. 7L) at 3h without changing the network burst rate (Fig. 7M) or network burst spike rate (Fig. 7N).

**Figure 7.**
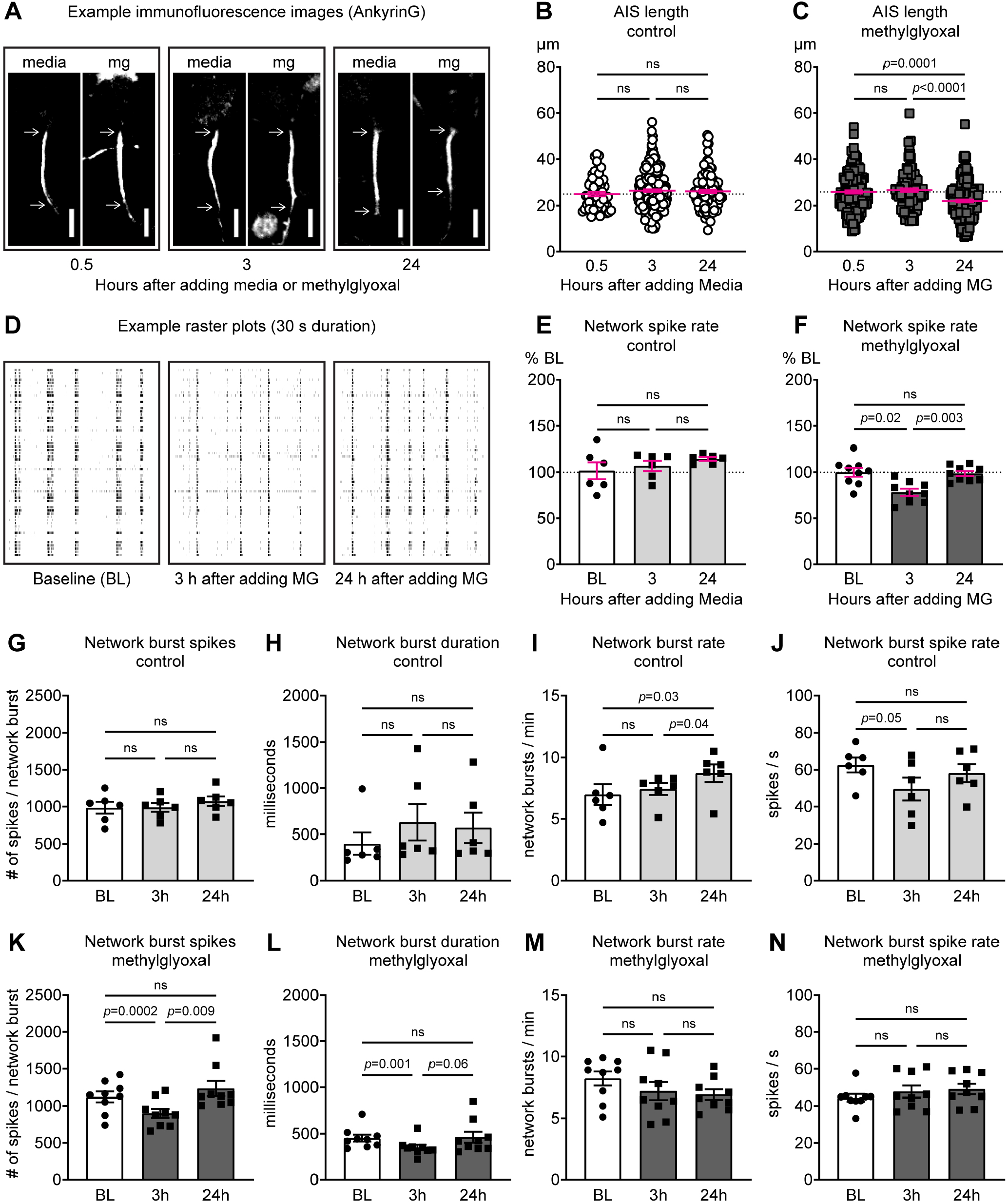
Time course of methylglyoxal-evoked AIS length and network activity changes. (A) Representative images of AnkyrinG (AIS) immunostaining at 0.5, 3, 24 h after media or methylglyoxal (mg) exposure. Scale bars = 10 µm. (B) AIS length at 0.5, 3, 24 h after media control treatment indicating no change in AIS length. (C) AIS length at 0.5, 3, 24 h after 100 µM methylglyoxal exposure indicating AIS shortening at 24 h. (D) Representative raster plots (30 s duration) showing spike activity (60 electrodes, one per row) during 15 min MEA recordings at baseline and 3 and 24 h after exposure to 100 µM methylglyoxal. (E) Network spike rate after media control exposure (n=6 MEAs from 2 preparations). (F) Network spike rate after methylglyoxal exposure (n=9 MEAs from 3 preparations). (G-J) Network spiking parameters after media control exposure. (K-N) Network spiking parameters after methylglyoxal exposure.

These time course analyses show that reduced network spiking by methylglyoxal returned to normal in parallel with the development of AIS shortening. In addition, they show that immediate, transient increase of spiking and bursting following methylglyoxal application (Fig. 6) did not lead to activity-dependent AIS shortening within 3 h (Fig. 7), even though methylglyoxal (300 µM) acutely depolarizes cortical neurons (de Arriba et al., 2006). This contrasts with rapid activity-dependent AIS shortening elicited by depolarizing high KCl within 1-3 h (Evans et al., 2015; Jamann et al., 2021). These observations hint that AIS shortening by methylglyoxal is likely different than homeostatic AIS shortening after depolarization by KCl.

### 3.8. Methylglyoxal shortens the AIS in multiple cell types

Previous studies indicate that AIS plasticity is cell-type dependent (Grubb and Burrone, 2010; Evans et al., 2013; Chand et al., 2015). To test if restoration of network activity (Fig. 7) is associated with AIS shortening in specific types of cells, we immunostained for MAP2 to label all neurons, βIV spectrin to label the AIS, and Ca^2+^/calmodulin-dependent protein kinase II alpha (CaMKIIα) to identify excitatory (CaMKIIα+) or putative inhibitory (CaMKIIα-) neurons. Representative images of postnatal cortical cultures at DIV14 double-labeled for MAP2 and CaMKIIα are shown (Fig. 8A). Exposure to 100 µM methylglyoxal for 24 h did not change the distribution of CaMKIIα+/- neurons: the mean percentage of all AIS measured that originated from CaMKIIα+ neurons was similar between media (79.7 ± 1.8 %, n=12 coverslips from 3 preparations) and methylglyoxal (77.3 ± 1.1 %, n=12 coverslips from 3 preparations) treatments (*p* = 0.45) (Fig. 8B). Analysis of only neurons treated with media control indicated that AIS length in CaMKIIα-neurons (27.81 µm) was ∼7.5% shorter than CaMKIIα+ neurons (30.08 µm). This shorter AIS length in inhibitory neurons is consistent with a previous study of hippocampal cultures from GAD67-GFP transgenic mice (Prestigio et al., 2019). Representative images of AIS length in CaMKIIα+ and CaMKIIα-neurons after 24 h exposure to 100 µM methylglyoxal are shown (Fig. 8C). In CaMKIIα+ neurons, AIS length after methylglyoxal exposure (26.25 ± 0.27 µm, n=296 AIS from 3 preparations) was shorter than media control (30.08 ± 0.30 µm, n=298 AIS from 3 preparations) (*p* < 0.0001; unpaired homoscedastic two-tailed t-test, 12.7 % reduction). Similarly, in CaMKIIα-neurons, AIS length after methylglyoxal exposure (23.65 ± 0.42 µm, n=86 AIS from 3 preparations) was shorter than media control (27.81 ± 0.47 µm, n=76 AIS from 3 preparations) (*p* < 0.0001, unpaired homoscedastic two-tailed t-test, 15.0 % reduction) (Fig. 8D). Our results using two complementary AIS markers, AnkyrinG (Figs. 4, 5C, 7A-C) and βIV spectrin (Fig. 8C,D), demonstrate that acute elevation of methylglyoxal leads to reversible AIS shortening without cell death in both excitatory and inhibitory neuron populations at a dose that disrupts cellular methylglyoxal metabolism and changes network activity.

**Figure 8.**
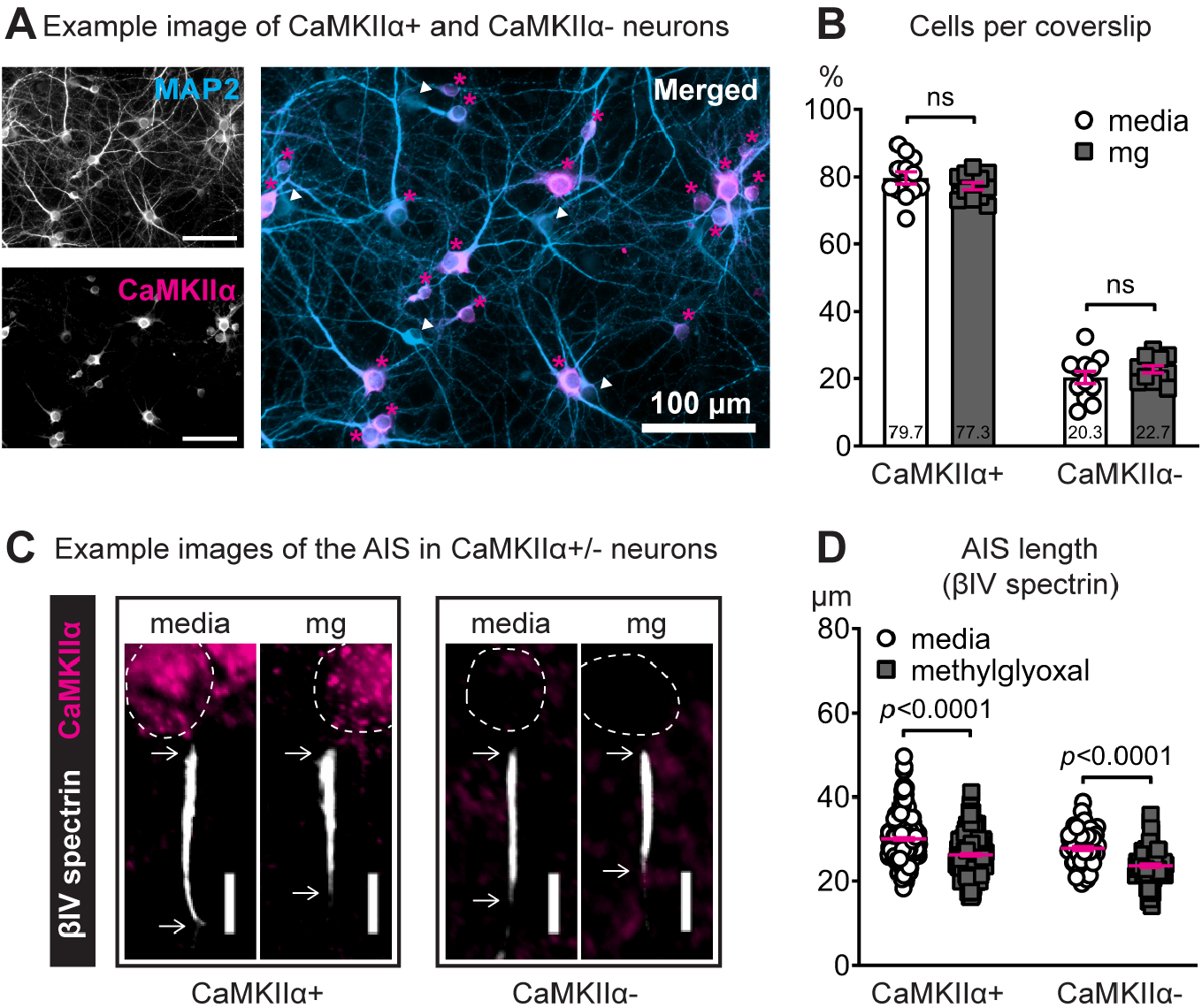
Methylglyoxal shortens the AIS in both excitatory and putative inhibitory neurons. (A) Representative images showing MAP2 (soma & dendrites) and CaMKIIα (excitatory neuron marker) immunostaining in DIV14 postnatal cortical cultures. CaMKIIα+ (magenta asterisks) and CaMKIIα-(white arrowheads) neurons are indicated in the merged image. Scale bar = 100 µm. (B) Quantification of the percentage of total cells counted that were CaMKIIα+ (excitatory) or CaMKIIα-(putative inhibitory) after 24 h exposure to 100 µM methylglyoxal (mg). Numbers within the bars indicate percentage of total cells counted. (C) Representative images of the AIS (βIV spectrin) in CaMKIIα+ and CaMKIIα-neurons after 24 h exposure to 100 µM methylglyoxal. Arrows denote the start and end of the AIS. Dashed lines indicate the neuronal soma. Scale bar = 10 µm. (D) Quantification of AIS length (βIV spectrin) after 24 h exposure to media or 100 μM methylglyoxal in CaMKIIα+ and CaMKIIα- neurons.

## 4. Discussion

### 4.1. Methylglyoxal mediates AIS shortening

Because of the association between AIS shortening and cognitive impairment in type 2 diabetes we sought to identify the initiator of AIS shortening. Canonical features of type 2 diabetes, insulin resistance (Fig. 1) or high glucose (Fig. 2), did not dramatically alter AIS length in mouse cortical neuron culture. In contrast, increase of methylglyoxal (100 µM) produced 11.5% AIS shortening (Fig. 4) that was recoverable and not associated with cell death (Fig. 5). This is consistent with 8-16% AIS shortening in the prefrontal cortex of type 2 diabetic db/db mice at 10 weeks of age with lack of apoptotic cell death (Yermakov et al., 2018). Methylglyoxal elevation leads to accumulation of MG-H1 in patients with type 2 diabetes (Schalkwijk and Stehouwer, 2020) and both methylglyoxal (Bierhaus et al., 2012) and MG-H1 (Griggs et al., 2019) are elevated in db/db mice. Adding methylglyoxal to culture media increased MG-H1 *in vitro* (Fig. 3), indicating that our culture model recapitulates disruption of methylglyoxal metabolism that might be relevant to diabetes. These results suggest that methylglyoxal is a key mediator of AIS shortening in the type 2 diabetic brain. However, we recognize the reductionist *in vitro* approach herein cannot completely recapitulate chronic diabetic conditions *in vivo*. Nevertheless, it is reasonable to think that prolonged increase of methylglyoxal, or the combined disruption of methylglyoxal, insulin, and glucose metabolism, in type 2 diabetes could result in sustained AIS shortening and CNS dysfunction.

### 4.2. AIS shortening may contribute to cognitive impairment and neuronal network dysfunction

Current and previous results suggest that methylglyoxal-evoked AIS shortening could be involved in the pathophysiology of diabetic brain complications. Methylglyoxal metabolism is disrupted in patients with diabetes (Sveen et al., 2013; Andersen et al., 2018) or cognitive deficits (Beeri et al., 2011; Srikanth et al., 2013; Cai et al., 2014). In animals, a single intracerebroventricular administration (Qi et al., 2017; Lissner et al., 2021) or repeated injection (Hansen et al., 2016; Szczepanik et al., 2020) of methylglyoxal induces cognitive impairment. These observations suggest that persistent elevation of methylglyoxal, such as in diabetes, could produce long-lasting changes in the operation of CNS networks *in vivo*. Consistent with disruption of neuronal network operation, we observed transient alterations of MEA activity after just a single bolus of methylglyoxal *in vitro* (Figs. 6,7). Importantly, methylglyoxal is increased in db/db mice (Bierhaus et al., 2012; Griggs et al., 2019) with cognitive impairment and shortened AIS in the prefrontal cortex and hippocampus (Yermakov et al., 2018, 2019), suggesting a connection between methylglyoxal, type 2 diabetes, AIS shortening, and cognitive impairment. Strengthening a mechanistic link between subtle changes in AIS geometry and CNS dysfunction, pharmacological treatment of nerve-injured mice simultaneously alleviated both AIS shortening and cognitive impairment (Shiers et al., 2018).

Little is known about the influence of AIS shortening on neuronal network operation and thus cognitive function. At the single-cell level, AIS shortening reduces neuronal excitability according to *in silico* (Baalman et al., 2013; Vascak et al., 2017; Goethals and Brette, 2020), *in vitro* (Evans et al., 2015), and *in vivo* (Jamann et al., 2021) reports. However, other studies show that AIS shortening does not reduce excitability when associated with changes in Nav phosphorylation (Evans et al., 2015), loss of Kv1 channels (Sanders et al., 2020), or alteration of action potential backpropagation (Radecki et al., 2018). At the network level, one study reported AIS shortening and reduced MEA spiking, but the cultures consisted of only excitatory neurons (Sohn et al., 2019). Here, we measured AIS geometry and MEA activity in neuronal networks containing both excitatory and inhibitory neurons. We observed AIS shortening in both CaMKII+ and CaMKII-neurons (Fig. 8) when network spiking returned to basal levels at 24 h, an increase from diminished activity at 3 h (Fig. 7). This seems counterintuitive if the current understanding is that AIS shortening reduces excitability. However, comparison of single neuron excitability versus MEA activity is difficult since many factors (e.g., composition of the neuronal network) can dictate network operation. We offer some speculative explanations. First, if inhibitory output plays a predominant role in network activity, then shortened AIS in inhibitory neurons would reduce the inhibitory drive onto excitatory neurons causing network activity to increase. This suggests diminished network activity when AIS shortening occurs in systems consisting of excitatory neurons only. Indeed, shorter AIS length associated with reduced MEA network burst rate in human iPSC-derived excitatory neuron cultures lacking inhibitory neurons (Sohn et al., 2019). Second, AIS shortening could reduce excitability in inhibitory neurons but not excitatory neurons since differences in the size or shape of these neurons can affect how AIS shortening modifies their excitability (Gulledge and Bravo, 2016). Third, altered Nav phosphorylation or Kv1 levels (discussed above) could differentially affect the functional outcome of AIS shortening in excitatory versus inhibitory neurons. Finally, spontaneous spiking does not reveal how underlying changes (i.e., AIS shortening) in the network might affect the response to subsequent perturbation. Thus, homeostatic capability of the network could still be impaired. This was previously reported for mutant tau excitatory neurons with shorter AIS, which led to a blunted response in MEA activity after challenge with depolarizing KCl (Sohn et al., 2019). Understanding how simultaneous AIS shortening in multiple cell types translates to the recovery of network function will be critical going forward.

### 4.3. How does methylglyoxal trigger network activity changes and AIS shortening?

AIS shortening has been described as a homeostatic mechanism that reduces neuronal excitability in response to depolarization or elevated activity (Evans et al., 2015; Jamann et al., 2021). In our study, AIS shortening could be a homeostatic response to the perturbation of network activity by methylglyoxal since we observed AIS shortening when activity was increasing back to baseline levels between 3 and 24 h (Fig. 7). However, the pattern of AIS shortening in the current results is different than homeostatic AIS shortening reported previously (Evans et al., 2015). Evans *et al* (2015) described cell-type specific AIS shortening in excitatory CA3 but not inhibitory GABAergic hippocampal neurons after 3 h exposure to KCl. We observed cell-type independent AIS shortening in both excitatory CaMKIIα+ and putative inhibitory CaMKIIα-cortical neurons after 24 h exposure to methylglyoxal (Fig. 8). Even though methylglyoxal depolarizes cortical neurons (de Arriba et al., 2006) and immediately increased MEA activity (Fig. 6), we did not observe homeostatic AIS shortening at 3 h. These observations hint that the mechanistic underpinnings of AIS shortening by methylglyoxal are different than homeostatic AIS shortening by depolarizing KCl.

If AIS shortening by methylglyoxal is not a mechanism of homeostatic plasticity, then how does network activity recover? AIS geometry is unlikely to be the sole determinant of network operation after methylglyoxal exposure. For example, homeostatic increase of synaptic input could lead to recovery of network activity despite AIS shortening. Indeed, methylglyoxal increases both evoked and spontaneous excitatory postsynaptic potentials in cockroach abdominal ganglia (Chambers et al., 1985), but whether it alters synaptic efficacy in mouse cortical neurons is unknown. Type 2 diabetic db/db mice show AIS shortening (Yermakov et al., 2018, 2019), but there are also reports of altered synaptic plasticity (Li et al., 2002; Stranahan et al., 2008), decreased dendritic spine density (Stranahan et al., 2009; Chen et al., 2014), and progressive cortical atrophy (Ramos-Rodriguez et al., 2013). In addition, elongation of the node of Ranvier after exposing myelinated CNS nerves to methylglyoxal *ex vivo* (Griggs et al., 2018) is similar to the elongated nodes present in older db/db mice (Yermakov et al., 2019). A better understanding of how concurrent changes at the AIS, node of Ranvier, and synapse affect network operation and cognitive function in neurological disease is needed.

The detailed molecular mechanisms mediating changes in network operation and AIS geometry in the current study remain unknown. Methylglyoxal may alter network activity by depolarization (de Arriba et al., 2006; Bierhaus et al., 2012), glutamate (de Arriba et al., 2006; Lissner et al., 2021), sodium channels (Bierhaus et al., 2012), the cation channel transient receptor potential ankyrin A (TRPA1) (Eberhardt et al., 2012; Andersson et al., 2013), GABA_A_ receptor (Distler et al., 2012), or mitochondrial ATP depletion (de Arriba et al., 2006, 2007). Methylglyoxal-evoked AIS shortening could involve calcium, calpain, calcineurin, or tau. Methylglyoxal promotes calcium mobilization in neurons (Radu et al., 2012; Griggs et al., 2019), activation of calpain (Griggs et al., 2018) and calcineurin (Maeta et al., 2005), and hyperphosphorylation of tau (Li et al., 2012). Likewise, calcium channels (Grubb and Burrone, 2010; Evans et al., 2013, 2015; Chand et al., 2015), calpain (Schafer et al., 2009; Clark et al., 2017), calcineurin (Evans et al., 2015), and tau (Sohn et al., 2019) are all implicated in AIS disruption.

### 4.4. Conclusion

Our results indicate that methylglyoxal is a key mediator of AIS shortening and network activity changes in cortical neurons. Further investigation of the mechanisms of methylglyoxal-evoked AIS shortening and perturbation of network operation could have a beneficial impact on a wide variety of conditions associated with disrupted methylglyoxal metabolism such as type 2 diabetes, Alzheimer’s disease, and aging.

## Acknowledgements

Funding sources R01NS107398 and R03NS112981. Mike Bottomley (Statistical Consulting Center, Wright State University) for help with experimental design and statistical analysis. Domenica Drouet and Dr. Andrew Koesters (Wright State University) for help with carrying out experiments and critical feedback. Dr. Thomas Fleming, Dr. Sebastian Brings, and Dr. Peter Nawroth (University of Heidelberg) for providing the Rat anti-MG-H1 1D2 tissue culture supernatant unpurified antibody and critical feedback. The lab of Dr. Peter Wenner (Emory University) for critical feedback.

## References

Andersen ST, Witte DR, Dalsgaard EM, Andersen H, Nawroth P, Fleming T, Jensen TSM, Finnerup NB, Jensen TSM, Lauritzen T, Feldman EL, Callaghan BC, Charles M (2018) Risk factors for incident diabetic polyneuropathy in a cohort with screen-detected type 2 diabetes followed for 13 years: Addition-Denmark. Diabetes Care 41:1068–1075.

Andersson DA, Gentry C, Light E, Vastani N, Vallortigara J, Bierhaus A, Fleming T, Bevan S (2013) Methylglyoxal Evokes Pain by Stimulating TRPA1. PLoS One 8:1–9.

Atapour N, Rosa MGP (2017) Age-related plasticity of the axon initial segment of cortical pyramidal cells in marmoset monkeys. Neurobiol Aging 57:95–103.

Baalman KL, Cotton RJ, Rasband SN, Rasband MN (2013) Blast wave exposure impairs memory and decreases axon initial segment length. J Neurotrauma 30:741–751.

Beaudoin GMJJ, Lee SH, Singh D, Yuan Y, Ng YG, Reichardt LF, Arikkath J (2012) Culturing pyramidal neurons from the early postnatal mouse hippocampus and cortex. Nat Protoc 7:1741– 1754.

Beeri MS, Moshier E, Schmeidler J, Godbold J, Uribarri J, Reddy S, Sano M, Grossman HT, Cai W, Vlassara H, Silverman JM (2011) Serum concentration of an inflammatory glycotoxin, methylglyoxal, is associated with increased cognitive decline in elderly individuals. Mech Ageing Dev 132:583–587.

Belanger M, Yang J, Petit JM, Laroche T, Magistretti PJ, Allaman I (2011) Role of the glyoxalase system in astrocyte-mediated neuroprotection. J Neurosci 31:18338–18352.

Bender KJ, Trussell LO (2012) The physiology of the axon initial segment. Annu Rev Neurosci 35:249–265.

Bierhaus A et al. (2012) Methylglyoxal modification of Nav1.8 facilitates nociceptive neuron firing and causes hyperalgesia in diabetic neuropathy. Nat Med 18:926–933.

Biessels GJ, Despa F (2018) Cognitive decline and dementia in diabetes mellitus: mechanisms and clinical implications. Nat Rev Endocrinol 14:591–604.

Biessels GJ, Strachan MWJ, Visseren FLJ, Kappelle LJ, Whitmer RA (2014) Dementia and cognitive decline in type 2 diabetes and prediabetic stages: Towards targeted interventions. Lancet Diabetes Endocrinol 2:246–255.

Buffington S, Rasband M (2011) The axon initial segment in nervous system disease and injury. Eur J Neurosci 34:1609–1619.

Cai W, Uribarri J, Zhu L, Chen X, Swamy S, Zhao Z, Grosjean F, Simonaro C, Kuchel GA, Schnaider-Beeri M, Woodward M, Striker GE, Vlassara H (2014) Oral glycotoxins are a modifiable cause of dementia and the metabolic syndrome in mice and humans. Proc Natl Acad Sci U S A 111:4940– 4945.

Chambers PL, Davies MG, Rowan MJ (1985) The Effects of Methylglyoxal on Central Synaptic Transmission in the Isolated Nerve Cord of the Cockroach (Periplaneta Americana L) BT -Receptors and Other Targets for Toxic Substances. In: Arch Toxicol (Chambers PL, Cholnoky E, Chambers CM, eds), pp 337–341. Berlin, Heidelberg: Springer Berlin Heidelberg.

Chand AN, Galliano E, Chesters RA, Grubb MS (2015) A Distinct Subtype of Dopaminergic Interneuron Displays Inverted Structural Plasticity at the Axon Initial Segment. J Neurosci 35:1573–1590.

Charkhkar H, Meyyappan S, Matveeva E, Moll JR, McHail DG, Peixoto N, Cliff RO, Pancrazio JJ (2015) Amyloid beta modulation of neuronal network activity in vitro. Brain Res 1629:1–9.

Chen J, Liang L, Zhan L, Zhou Y, Zheng L, Sun X, Gong J, Sui H, Jiang R, Zhang F, Zhang L (2014) ZiBuPiYin recipe protects db/db mice from diabetes-associated cognitive decline through improving multiple pathological changes. PLoS One 9:1–12.

Clark KC, Sword BA, Dupree JL (2017) Oxidative Stress Induces Disruption of the Axon Initial Segment. ASN Neuro 9:175909141774542.

de Arriba SG et al. (2006) Carbonyl stress and NMDA receptor activation contribute to methylglyoxal neurotoxicity. Free Radic Biol Med 40:779–790.

de Arriba SG, Stuchbury G, Yarin J, Burnell J, Loske C, Münch G (2007) Methylglyoxal impairs glucose metabolism and leads to energy depletion in neuronal cells-protection by carbonyl scavengers. Neurobiol Aging 28:1044–1050.

Ding Y, Chen T, Wang Q, Yuan Y, Hua T (2018) Axon initial segment plasticity accompanies enhanced excitation of visual cortical neurons in aged rats. Neuroreport 29:1537–1543.

Distler MG, Plant LD, Sokoloff G, Hawk AJ, Aneas I, Wuenschell GE, Termini J, Meredith SC, Nobrega MA, Palmer AA (2012) Glyoxalase 1 increases anxiety by reducing GABA A receptor agonist methylglyoxal. J Clin Invest 122:2306–2315.

Düll MM, Riegel K, Tappenbeck J, Ries V, Strupf M, Fleming T, Sauer SK, Namer B (2019) Methylglyoxal causes pain and hyperalgesia in human through C-fiber activation. Pain 160:2497– 2507.

Eberhardt MJ, Filipovic MR, Leffler A, De La Roche J, Kistner K, Fischer MJ, Fleming T, Zimmermann K, Ivanovic-Burmazovic I, Nawroth PP, Bierhaus A, Reeh PW, Sauer SK (2012) Methylglyoxal activates nociceptors through transient receptor potential channel A1 (TRPA1): A possible mechanism of metabolic neuropathies. J Biol Chem 287:28291–28306.

Evans MD, Dumitrescu AS, Kruijssen DLH, Taylor SE, Grubb MS (2015) Rapid Modulation of Axon Initial Segment Length Influences Repetitive Spike Firing. Cell Rep 13:1233–1245.

Evans MD, Sammons RP, Lebron S, Dumitrescu AS, Watkins TBK, Uebele VN, Renger JJ, Grubb MS (2013) Calcineurin Signaling Mediates Activity-Dependent Relocation of the Axon Initial Segment. J Neurosci 33:6950–6963.

Evans MD, Tufo C, Dumitrescu AS, Grubb MS (2017) Myosin II activity is required for structural plasticity at the axon initial segment. Eur J Neurosci:1–7.

Fong M, Newman JP, Potter SM, Wenner P (2015) Upward synaptic scaling is dependent on neurotransmission rather than spiking. Nat Commun 6:1–11.

Fruscione F, Valente P, Sterlini B, Romei A, Baldassari S, Fadda M, Prestigio C, Giansante G, Sartorelli J, Rossi P, Rubio A, Gambardella A, Nieus T, Broccoli V, Fassio A, Baldelli P, Corradi A, Zara F, Benfenati F (2018) PRRT2 controls neuronal excitability by negatively modulating Na+ channel 1.2/1.6 activity. Brain 141:1000–1016.

Galiano MR, Jha S, Ho TSY, Zhang C, Ogawa Y, Chang KJ, Stankewich MC, Mohler PJ, Rasband MN (2012) A distal axonal cytoskeleton forms an intra-axonal boundary that controls axon initial segment assembly. Cell 149:1125–1139.

Goethals S, Brette R (2020) Theoretical relation between axon initial segment geometry and excitability.Elife 9:834671.

Griggs RB, Donahue RR, Adkins BG, Anderson KL, Thibault O, Taylor BK (2016) Pioglitazone inhibits the development of hyperalgesia and sensitization of spinal nociresponsive neurons in type 2 diabetes. J Pain 17:359–373.

Griggs RB, Santos DF, Laird DE, Doolen S, Donahue RR, Wessel CR, Fu W, Sinha GP, Wang P, Zhou J, Brings S, Fleming T, Nawroth PP, Susuki K, Taylor BK (2019) Methylglyoxal and a spinal TRPA1-AC1-Epac cascade facilitate pain in the db/db mouse model of type 2 diabetes. Neurobiol Dis 127:76–86.

Griggs RB, Yermakov LM, Drouet DE, Nguyen DVM, Susuki K (2018) Methylglyoxal Disrupts Paranodal Axoglial Junctions via Calpain Activation. ASN Neuro 10:175909141876617.

Grubb MS, Burrone J (2010) Activity-dependent relocation of the axon initial segment fine-tunes neuronal excitability. Nature 465:1070–1074.

Gulledge AT, Bravo JJ (2016) Neuron morphology influences axon initial segment plasticity. eNeuro 3:255–265.

Guo Y, Su ZJ, Chen YK, Chai Z (2017) Brain-derived neurotrophic factor/neurotrophin 3 regulate axon initial segment location and affect neuronal excitability in cultured hippocampal neurons. J Neurochem 142:260–271.

Haddad M, Perrotte M, Khedher MR Ben Demongin C, Lepage A, Fülöp T, Ramassamy C (2019) Methylglyoxal and glyoxal as potential peripheral markers for mci diagnosis and their effects on the expression of neurotrophic, inflammatory and neurodegenerative factors in neurons and in neuronal derived-extracellular vesicles. Int J Mol Sci 20:4906.

Hales CM, Rolston JD, Potter SM (2010) How to culture, record and stimulate neuronal networks on micro-electrode arrays (MEAs). J Vis Exp:1–7.

Hamada MS, Kole MHP (2015) Myelin Loss and Axonal Ion Channel Adaptations Associated with Gray Matter Neuronal Hyperexcitability. J Neurosci 35:7272–7286.

Hansen F, Pandolfo P, Galland F, Torres FV, Dutra MF, Batassini C, Guerra MC, Leite MC, Gonçalves CA (2016) Methylglyoxal can mediate behavioral and neurochemical alterations in rat brain. Physiol Behav 164:93–101.

Harty RC, Kim TH, Thomas EA, Cardamone L, Jones NC, Petrou S, Wimmer VC (2013) Axon initial segment structural plasticity in animal models of genetic and acquired epilepsy. Epilepsy Res 105:272–279.

Hinman JD, Rasband MN, Carmichael ST (2013) Remodeling of the axon initial segment after focal cortical and white matter stroke. Stroke 44:182–189.

Huang CY, Rasband MN (2018) Axon initial segments: structure, function, and disease. Ann N Y Acad Sci 1420:1–16.

Huang Q, Chen Y, Gong N, Wang YX (2016) Methylglyoxal mediates streptozotocin-induced diabetic neuropathic pain via activation of the peripheral TRPA1 and Nav1.8 channels. Metabolism 65:463– 474.

Irshad Z, Xue M, Ashour A, Larkin JR, Thornalley PJ, Rabbani N (2019) Activation of the unfolded protein response in high glucose treated endothelial cells is mediated by methylglyoxal. Sci Rep 9:7889.

Jamann N, Dannehl D, Lehmann N, Wagener R, Thielemann C, Schultz C, Staiger J, Kole MHP, Engelhardt M (2021) Sensory input drives rapid homeostatic scaling of the axon initial segment in mouse barrel cortex. Nat Commun 12:1–14.

Jamann N, Jordan M, Engelhardt M (2018) Activity-dependent axonal plasticity in sensory systems. Neuroscience 368:268–282.

Kim B, Sullivan KA, Backus C, Feldman EL (2011) Cortical Neurons Develop Insulin Resistance and Blunted Akt Signaling: A Potential Mechanism Contributing to Enhanced Ischemic Injury in Diabetes. Antioxid Redox Signal 14:1829–1839.

Kuhla B, Boeck K, Lüth H-J, Schmidt A, Weigle B, Schmitz M, Ogunlade V, Münch G, Arendt T (2006) Age-dependent changes of glyoxalase I expression in human brain. Neurobiol Aging 27:815–822.

Kuhla B, Boeck K, Schmidt A, Ogunlade V, Arendt T, Münch G, Lüth HJ (2007) Age- and stage-dependent glyoxalase I expression and its activity in normal and Alzheimer’s disease brains. Neurobiol Aging 28:29–41.

Kuhla B, Lüth HJ, Haferburg D, Boeck K, Arendt T, Münch G (2005) Methylglyoxal, glyoxal, and their detoxification in Alzheimer’s disease. Ann N Y Acad Sci 1043:211–216.

Kurz A, Rabbani N, Walter M, Bonin M, Thornalley P, Auburger G, Gispert S (2011) Alpha-synuclein deficiency leads to increased glyoxalase i expression and glycation stress. Cell Mol Life Sci 68:721–733.

Li XH, Xie JZ, Jiang X, Lv BL, Cheng XS, D. LL, Zhang JY, Wang JZ, Zhou XW (2012) Methylglyoxal induces tau hyperphosphorylation via promoting ages formation. NeuroMolecular Med 14:338–348.

Li XL, Aou S, Oomura Y, Hori N, Fukunaga K, Hori T (2002) Impairment of long-term potentiation and spatial memory in leptin receptor-deficient rodents. Neuroscience 113:607–615.

Lissner LJ, Rodrigues L, Wartchow KM, Borba E, Bobermin LD, Fontella FU, Hansen F, Quincozes-Santos A, Souza DOG, Gonçalves CA (2021) Short-Term Alterations in Behavior and Astroglial Function After Intracerebroventricular Infusion of Methylglyoxal in Rats. Neurochem Res 46:183– 196.

Liu C-CC, Zhang X-SS, Ruan Y-TT, Huang Z-XX, Zhang S-BB, Liu M, Luo H-JJ, Wu S-LL, Ma C (2017) Accumulation of methylglyoxal increases the advanced glycation end products levels in DRG and contributes to lumbar disc herniation-induced persistent pain. J Neurophysiol 118:jn.00745.2016.

Maeta K, Izawa S, Inoue Y (2005) Methylglyoxal, a metabolite derived from glycolysis, functions as a signal initiator of the high osmolarity glycerol-mitogen-activated protein kinase cascade and calcineurin/Crz1-mediated pathway in Saccharomyces cerevisiae. J Biol Chem 280:253–260.

Marin MA, Ziburkus J, Jankowsky J, Rasband MN (2016) Amyloid-β plaques disrupt axon initial segments. Exp Neurol 281:93–98.

Nelson AD, Jenkins PM (2017) Axonal Membranes and Their Domains: Assembly and Function of the Axon Initial Segment and Node of Ranvier. Front Cell Neurosci 11:1–17.

Nishimoto S, Koike S, Inoue N, Suzuki T, Ogasawara Y (2017) Activation of Nrf2 attenuates carbonyl stress induced by methylglyoxal in human neuroblastoma cells?: Increase in GSH levels is a critical event for the detoxification mechanism. Biochem Biophys Res Commun 483:1–6.

Percie du Sert N et al. (2020) Reporting animal research: Explanation and elaboration for the ARRIVE guidelines 2.0. PLOS Biol 18:e3000411.

Potter SM, DeMarse TB (2001) A new approach to neural cell culture for long-term studies. J Neurosci Methods 110:17–24.

Prestigio C, Ferrante D, Valente P, Casagrande S, Albanesi E, Yanagawa Y, Benfenati F, Baldelli P (2019) Spike-Related Electrophysiological Identification of Cultured Hippocampal Excitatory and Inhibitory Neurons. Mol Neurobiol 56:6276–6292.

Qi LQ, Chen Z, Wang YP, Liu XY, Liu XH, Ke LF, Zheng Z, Lin XW, Zhou Y, Wu LJ, Liu L Bin (2017) Subcutaneous liraglutide ameliorates methylglyoxal-induced alzheimer-like tau pathology and cognitive impairment by modulating tau hyperphosphorylation and glycogen synthase kinase-3β. Am J Transl Res 9:247–260.

Rabbani N, Thornalley PJ (2012) Methylglyoxal, glyoxalase 1 and the dicarbonyl proteome. Amino Acids 42:1133–1142.

Radecki DZ, Johnson EL, Brown AK, Meshkin NT, Perrine SA, Gow A (2018) Corticohippocampal Dysfunction In The OBiden Mouse Model Of Primary Oligodendrogliopathy. Sci Rep 8:1–18.

Radu BM, Dumitrescu DI, Mustaciosu CC, Radu M (2012) Dual effect of methylglyoxal on the intracellular Ca2+ signaling and neurite outgrowth in mouse sensory neurons. Cell Mol Neurobiol 32:1047–1057.

Ramos-Rodriguez JJ, Ortiz O, Jimenez-Palomares M, Kay KR, Berrocoso E, Murillo-Carretero MI, Perdomo G, Spires-Jones T, Cozar-Castellano I, Lechuga-Sancho AM, Garcia-Alloza M (2013) Differential central pathology and cognitive impairment in pre-diabetic and diabetic mice. Psychoneuroendocrinology 38:2462–2475.

Rasband MN (2010) The axon initial segment and the maintenance of neuronal polarity. Nat Rev Neurosci 11:552–562.

Sanders SS, Hernandez LM, Soh H, Karnam S, Walikonis RS, Tzingounis A V, Thomas GM (2020) The palmitoyl acyltransferase ZDHHC14 controls Kv1-family potassium channel clustering at the axon initial segment. Elife 9.

Schafer DP, Jha S, Liu F, Akella T, McCullough LD, Rasband MN (2009) Disruption of the Axon Initial Segment Cytoskeleton Is a New Mechanism for Neuronal Injury. J Neurosci 29:13242–13254.

Schalkwijk CG, Stehouwer CDA (2020) Methylglyoxal, a highly reactive dicarbonyl compound, in diabetes, its vascular complications, and other age-related diseases. Physiol Rev 100:407–461.

Shiers S, Pradhan G, Mwirigi J, Mejia G, Ahmad A, Kroener S, Price T (2018) Neuropathic pain creates an enduring prefrontal cortex dysfunction corrected by the type II diabetic drug metformin but not by gabapentin. J Neurosci 38:7337–7350.

Sohn PD, Huang CT-L, Yan R, Fan L, Tracy TE, Camargo CM, Montgomery KM, Arhar T, Mok S-A, Freilich R, Baik J, He M, Gong S, Roberson ED, Karch CM, Gestwicki JE, Xu K, Kosik KS, Gan L (2019) Pathogenic Tau Impairs Axon Initial Segment Plasticity and Excitability Homeostasis. Neuron 104:458-470.e5.

Sri S, Pegasiou CM, Cave CA, Hough K, Wood N, Gomez-Nicola D, Deinhardt K, Bannerman D, Perry VH, Vargas-Caballero M (2019) Emergence of synaptic and cognitive impairment in a mature-onset APP mouse model of Alzheimer’s disease. Acta Neuropathol Commun 7:25.

Srikanth V, Westcott B, Forbes J, Phan TG, Beare R, Venn A, Pearson S, Greenaway T, Parameswaran V, Münch G (2013) Methylglyoxal, cognitive function and cerebral atrophy in older people. Journals Gerontol -Ser A Biol Sci Med Sci 68:68–73.

Stranahan AM, Arumugam T V., Cutler RG, Lee K, Egan JM, Mattson MP (2008) Diabetes impairs hippocampal function through glucocorticoid-mediated effects on new and mature neurons. Nat Neurosci 11:309–317.

Stranahan AM, Lee K, Martin B, Maudsley S, Golden E, Cutler RG, Mattson MP (2009) Voluntary exercise and caloric restriction enhance hippocampal dendritic spine density and BDNF levels in diabetic mice. Hippocampus 19:951–961.

Sveen KA, Karimé B, Jørum E, Mellgren SI, Fagerland MW, Monnier VM, Dahl-Jørgensen K, Hanssen KF (2013) Small-and large-fiber neuropathy after 40 years of type 1 diabetes: Associations with glycem ic control andadvanced protein glycation: The Oslo Study. Diabetes Care 36:3712–3717.

Szczepanik JC, de Almeida GRL, Cunha MP, Dafre AL (2020) Repeated Methylglyoxal Treatment Depletes Dopamine in the Prefrontal Cortex, and Causes Memory Impairment and Depressive-Like Behavior in Mice. Neurochem Res 45:354–370.

Vascak M, Sun J, Baer M, Jacobs KM, Povlishock JT (2017) Mild traumatic brain injury evokes pyramidal neuron axon initial segment plasticity and diffuse presynaptic inhibitory terminal loss. Front Cell Neurosci 11:157.

Wagenaar DA, Pine J, Potter SM (2006) An extremely rich repertoire of bursting patterns during the development of cortical cultures. BMC Neurosci 7:11.

Wetzels S, Vanmierlo T, Scheijen JLJM, Van Horssen J, Amor S, Somers V, Schalkwijk CG, Hendriks JJA, Wouters K (2019) Methylglyoxal-derived advanced glycation endproducts accumulate in multiple sclerosis lesions. Front Immunol 10:1–9.

Yamada R, Kuba H (2016) Structural and Functional Plasticity at the Axon Initial Segment. Front Cell Neurosci 10:1–7.

Yang Y, Ogawa Y, Hedstrom KL, Rasband MN (2007) betaIV spectrin is recruited to axon initial segments and nodes of Ranvier by ankyrinG. J Cell Biol 176:509–519.

Yau KW, VanBeuningen SFB, Cunha-Ferreira I, Cloin BMC, VanBattum EY, Will L, Schätzle P, Tas RP, VanKrugten J, Katrukha EA, Jiang K, Wulf PS, Mikhaylova M, Harterink M, Pasterkamp RJ, Akhmanova A, Kapitein LC, Hoogenraad CC (2014) Microtubule minus-end binding protein CAMSAP2 controls axon specification and dendrite development. Neuron 82:1058–1073.

Yermakov LM, Drouet DE, Griggs RB, Elased KM, Susuki K (2018) Type 2 Diabetes Leads to Axon Initial Segment Shortening in db/db Mice. Front Cell Neurosci 12:1–13.

Yermakov LM, Griggs RB, Drouet DE, Sugimoto C, Williams MT, Vorhees C V., Susuki K (2019) Impairment of cognitive flexibility in type 2 diabetic db/db mice. Behav Brain Res 371:111978.

Zhang W, Bonadiman A, Ciorraga M, Benitez MJ, Garrido JJ (2019) P2Y1 purinergic receptor modulate axon initial segment initial development. Front Cell Neurosci 13:1–15.

